# Arl2 Associates with Cdk5rap2 to Regulate Cortical Development via Microtubule Organization

**DOI:** 10.1101/2024.01.16.575964

**Authors:** Dong-Liang Ma, Kun-Yang Lin, Suresh Divya, Jiaen Lin, Mahekta R. Gujar, Htet Yamin Aung, Ye Sing Tan, Yang Gao, Anselm S. Vincent, Teng Chen, Hongyan Wang

**Affiliations:** Program in Neuroscience and Behavioural Disorders, Duke-NUS Medical School, Singapore 169857; College of Forensic Medicine, Xi’an Jiaotong University Health Science Center, Xi’an, Shaanxi, PR China, 710061; The Key Laboratory of Health Ministry for Forensic Science, Xi’an Jiaotong University, Shaanxi, PR China, 710061; Department of Physiology, Yong Loo Lin School of Medicine, National University of Singapore, Singapore 117597; Integrative Sciences and Engineering Programme, National University of Singapore, 28 Medical Drive, Singapore 117456

**Keywords:** Arl2, neural stem cells, cortical development, neurogenesis, microtubules, Cdk5rap2/Cep215

## Abstract

ADP ribosylation factor-like GTPase 2 (Arl2) is crucial for controlling mitochondrial fusion and microtubule assembly in various organisms. Arl2 regulates the asymmetric division of neural stem cells in *Drosophila* via microtubule growth. However, the function of mammalian Arl2 during cortical development was unknown. Here, we demonstrate that mouse Arl2 plays a new role in corticogenesis via regulating microtubule growth, but not mitochondria functions. Arl2 knockdown leads to impaired proliferation of neural progenitor cells (NPCs) and neuronal migration. Arl2 knockdown in mouse NPCs significantly diminishes centrosomal microtubule growth and delocalization of centrosomal proteins Cdk5rap2 and γ-tubulin. Moreover, Arl2 physically associates with Cdk5rap2 by *in silico* prediction using AlphaFold Multimer and *in vitro* binding assays. Remarkably, Cdk5rap2 overexpression significantly rescues the neurogenesis defects caused by Arl2 knockdown. Therefore, Arl2 plays an important role in mouse cortical development through microtubule growth via the centrosomal protein Cdk5rap2.

## Introduction

Neural stem cells (NSCs) play a central role in the development of the mammalian brain. Cortical NSCs reside in the ventricular zone (VZ) and subventricular zone (SVZ), namely neuroepithelial cells and the apical radial glial cells, self-renew and proliferate to generate neurons that migrate to the cortical plate (CP) [1-8]. Both types of cells are collectively termed as neural stem and progenitor cells, hence referred to as neural progenitor cells (NPCs). NPCs divide either symmetrically or asymmetrically [1, 2]. Symmetric division of NPCs expands the stem cell pool during early neurogenesis [1, 2]. Subsequently, NPCs divide asymmetrically to generate intermediate progenitor cells that divide once to produce two neurons [1, 9]. The balance between the proliferation and differentiation of NPCs has a direct impact on neuron formation. Moreover, defects in NPC proliferation are associated with neurodevelopmental disorders [10-12].

Centrosomal proteins play crucial roles during mouse cortical development. The centrosome, composed of a pair of centrioles surrounded by pericentriolar material protein (PCM), is the major microtubule-organizing center (MTOC) that contributes to the formation of the mitotic spindle during cell division. A few centrosomal proteins including PCM1 and Cep120 play critical roles in brain development and variants in these two genes are associated with primary microcephaly [13-15]. CDK5 Regulatory Subunit Associated Protein 2 (Cdk5rap2/Cep215) is an evolutionarily conserved PCM protein that plays a crucial role in centrosomal duplication and maturation as well as microtubule organization in various organisms. Cdk5rap2 is critical for proliferation and differentiation of neuronal progenitor cells during mouse cortical development [16, 17]. Mutations in Cdk5rap2 are associated with congenital diseases such as primary microcephaly and primordial dwarfism [18, 19].

Arl2 (ADP-ribosylation factor-like 2) is an evolutionarily conserved small GTPase that is crucial for the formation of microtubules and maintaining centrosome integrity [20, 21]. Arl2 cycles between an inactive GDP-bound and an activated GTP-bound state and is a regulator of tubulin folding and microtubule biogenesis [9, 20-24]. Yeast orthologue of Arl2, together with TBCD and TBCE, forms a tubulin chaperone for microtubule biogenesis [25]. We previously showed that *Drosophila* Arl2 is essential for NSC polarity and microtubule growth [22]. Arl2 also plays a role in mitochondrial dynamics and function [24]. Arl2 regulates mitochondrial fusion when it is in the intermembrane space [26]. Arl2 also interacts with mitochondrial outer membrane proteins Miro1 and Miro2 to modulate mitochondrial transport and distribution [27]. Variants in human ARL2 and ARL2BP have been identified in eye disorders namely MRCS (microcornea, rod-cone dystrophy, cataract, and posterior staphyloma) syndrome and retinitis pigmentosa, respectively [28, 29].

Mammalian Arl2 is widely expressed in various tissues and is most abundant in the brain [30]. However, the role of mammalian Arl2 during brain development has not been established. In this study, we demonstrate a novel role for the mammalian Arl2 in cortical development. We show that Arl2 is required for the proliferation, migration and differentiation of mouse forebrain NPCs *in vitro* and *in vivo* by regulating centrosome assembly and microtubule growth in NPCs. Moreover, Arl2 co-localizes with Cdk5rap2 at the centrosomes and can physically associate with it. Finally, Arl2 functions upstream of Cdk5rap2 in regulating NPC proliferation and migration during mouse cortical development.

## RESULTS

### Arl2 knockdown results in a reduction in mNPC proliferation and neuronal migration

To examine the role of mouse Arl2 in cortical development, we silenced endogenous Arl2 expression in the primary culture of mouse neural progenitor cells (mNPCs) isolated from E14 mouse cortex *in vitro* (Supplementary Figure 1A) as well as in utero electroporated cells in mouse brain *in vivo* (Figure 1A) using short hairpin RNA (shRNA). We identified two independent shArl2-1 and shArl2-2 tagged with green fluorescent protein (GFP) which are capable of knocking down endogenous Arl2 expression. Upon Arl2 knock down (KD), Arl2 protein detected by anti-Arl2 antibodies in the Western blot (WB) was reduced to 42.52% (shArl2-1) and 20.37% (shArl2-2), respectively, compared with the control. Since knockdown by shArl2-2 is more efficient (Supplementary Figure 1B-C), we used shArl2-2 for the subsequent phenotypic analysis.

**Fig. 1.**
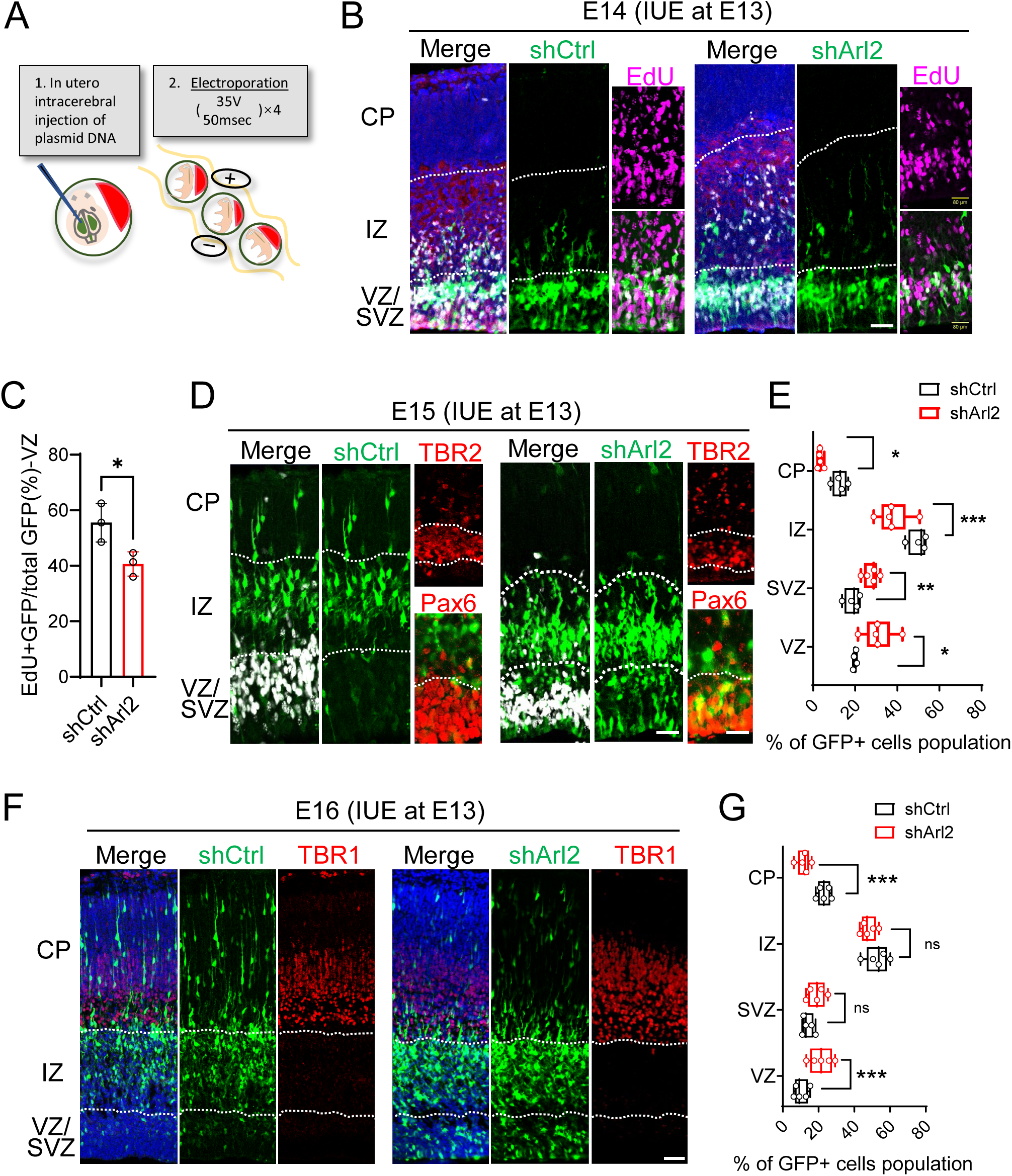
Arl2 knockdown results in a reduction in mNPC proliferation and neuronal migration. (A) Schematic representation of in-utero electroporation (IUE). (B) EdU labelling by ClickiT® EdU Imaging Kits. Brain slices from shCtrl (scrambled control) and shArl2 (Arl2 shRNA) groups at E14, one day after IUE, were labelled with EdU and GFP. (C) Bar graph showing reduced EdU incorporation upon Arl2 knockdown (40.66 ± 4.35% in shArl2 vs 55.54 ± 6.96% in shCtrl). The values represent the mean ± s.d. (n = 3 embryos). Student’s t-test, differences were considered significant at *p<0.05. (D) Cortical brain sections from shCtrl and shArl2 groups at E15, two days after IUE, were labelled with TBR2 (intermediate neural progenitor marker and labelling SVZ) or Pax6 (radial glia marker and labelling VZ) with GFP. (E) Box plots representing GFP+ cells in CP (shctrl: 12.68 ± 3.47%, shArl2: 3.42 ± 1.17%), IZ (shctrl: 49.99 ± 4.42%, shArl2: 38.00 ± 7.91%), SVZ (shctrl: 18.76 ± 3.71%, shArl2: 27.64 ± 3.53%) & VZ (shctrl: 19.92 ± 0.87%, shArl2: 30.94 ± 7.51%) showing defects in neuronal migration to CP upon Arl2 Knockdown compared to the control. The values represent the mean ± s.d. (shCtrl, n = 4 embryos; shArl2, n = 5 embryos). Multiple unpaired t tests, differences were considered significant at **p<0.05, ***p<0.001 and ****p<0.0001. (F) Cortical brain sections for shCtrl and shArl2 groups at E16, three days after IUE, were immunolabelled with TBR1 (immature neuron marker and labelling CP) and GFP. (G) Box plots representing GFP+ cells in CP (shctrl: 23.07 ± 3.61%, shArl2: 11.75 ± 3.67%,), IZ (shctrl: 52.82 ± 6.31%, shArl2: 48.00 ± 4.24%,), SVZ (shctrl: 13.94 ± 3.15%, shArl2: 18.82 ± 4.90%,) & VZ (shctrl: 10.17 ± 3.85%, shArl2: 21.42 ± 6.38%,) showing defects in neuronal migration to CP upon Arl2 Knockdown compared to the control. The values represent the mean ± s.d. (shCtrl, n = 5 embryos; shArl2, n = 5 embryos). Multiple unpaired t tests, differences were considered significant at **p<0.05 and ****p<0.0001. ns = non-significance. Scale bars; B and D = 50 µm, F = 80 µm.

To determine whether Arl2 KD affects the proliferation and differentiation of mNPCs during mouse cortical development, we silenced endogenous Arl2 expression in the primary culture of mNPCs *in vitro* by lentivirus (pPurGreen) infection in 48 h culture and pulse-labelled with EdU for 3 h before harvesting the cells. Remarkably, we observed a significant decrease in the proportion of EdU+ cells to 34.35% and Ki67+ cells to 36.98% in the shArl2 group as compared to the control group (46.03% and 60.09%, respectively) (Supplementary Figure 1D-F). Consistent with these observations, there was a significant reduction in cell proliferation upon Arl2 KD in mNPCs as compared to control in automated live-imaging analysis by Incucyte (Supplementary Figure 1G-H; Movie S1-2). Taken together, our data suggests that Arl2 is required for the proliferation of mNPCs *in vitro*.

To assess the impact of Arl2 knockdown (KD) on the proliferation, differentiation, and migration of mouse neural progenitor cells (mNPCs) during cortical development *in vivo*, we introduced Arl2 shRNA-2 via microinjection into the lateral ventricle of mouse embryos, followed by *in utero* electroporation (IUE) at embryonic day 13 (E13) (Figure 1A), and examined cortical neurogenesis from E14 to E16. At E14, one day after IUE and following 6-hour pulse-labelling with EdU before sample collection, the majority of GFP-positive cells were located in the ventricular zone (VZ) and subventricular zone (SVZ), with the minority population of cells migrating into the intermediate zone (IZ) in both control and shArl2 group (Figure 1B). Interestingly, in the VZ and SVZ, we observed a substantial reduction in the proportion of EdU+/GFP+ double-labelled cells in the shArl2 group (40.66%) as compared to the control group (55.54%) (Figure 1C). This data strongly suggests a notable impairment in the proliferation of NSCs due to Arl2 KD *in vivo*. At E15, majority of control GFP-positive (GFP+) cells were located in the IZ (49.99%) or had migrated into the cortical plate (CP) (12.68%), with a few GFP+ remaining in the VZ (19.92%) and SVZ (18.76%) (Figure 1D, E). In contrast, Arl2 knockdown resulted in significantly more cells remaining in the VZ (30.94%), SVZ (27.64%), and IZ (38.00%), with fewer cells at the CP (3.42%) as compared to the control group (Figure 1D, E). At E16, three days after IUE, more GFP+ cells were located in the IZ (52.82%) and CP (23.07%), while the rest of the cells persisted in the VZ (10.17%) and SVZ (13.94%) in the control group (Figure 1F, G). Remarkably, Arl2 knockdown caused a notable retention of cells in the VZ (21.42%), SVZ (18.82%), and IZ (48.00%), accompanied by much fewer cells at the CP (11.75%) (Figure 1F, G), suggesting a defect in neuronal migration.

In summary, our findings collectively highlight the critical role of Arl2 in the proliferation of NSCs and the migration of neuronal cells during cortical development.

### Overexpression mArl2 or hArl2 results in an increase in neuronal migration

Since the Arl2 KD led to decreased proliferation and migration of neuronal cells, we wondered whether overexpressing Arl2 had any effect on cortical development. We overexpressed both the human and mouse forms of wildtype Arl2 (FUtdTW-Arl2^WT^) driven by UbC promotor via microinjection into the lateral ventricle of mouse embryos, followed by IUE at embryonic day 13 (E13) (Figure 2A and C). Interestingly, at E16, 3 days after IUE, there was a significant increase in neuronal cell migration (tdTomato-positive (Td+) cells) to the CP in human Arl2^WT^ (hArl2^WT^) (VZ = 11.13%, SVZ = 18.21%, IZ = 40.25%, CP = 30.41%) as compared to control (VZ = 13.23%, SVZ = 19.47%, IZ = 45.92%, CP = 21.39%). Likewise, overexpression of mouse Arl2^WT^ (mArl2^WT^) also resulted in a significant increase in neuronal migration (Figure 2A and C, VZ = 13.92%, SVZ = 20.69%, IZ = 36.09%, CP = 29.30%) as compared to control, suggesting human and mouse forms Arl2 have conserved functions in neuronal migration (Figure 2A and C). Since human and mouse Arl2 show 96% homology and our overexpression results show similar phenotypes for both species, we used the human Arl2 for all subsequent overexpression experiments unless otherwise stated.

**Fig 2.**
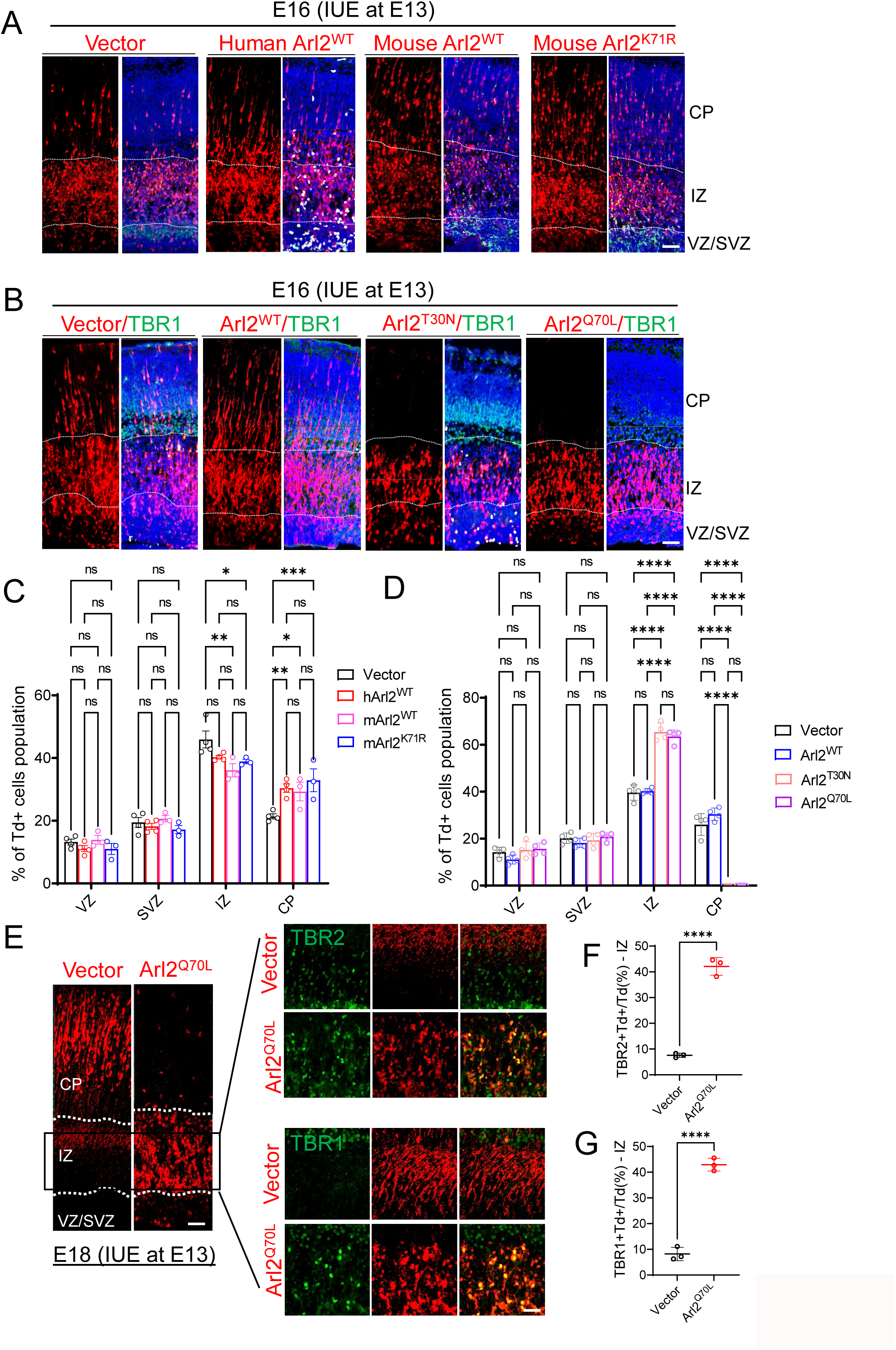
Overexpression various forms of Arl2 alters neuronal migration. (A) Cortical brain sections following overexpression of human Arl2^WT^ (hArl2^WT^), mouse Arl2^WT^ (mArl2^WT^) and mouse Arl2^K71R^ at E16, three days after IUE, were labelled with tdTomato (Td). (B) Cortical brain sections following overexpression of Arl2^WT^, the dominant-negative form (Arl2^T30N^) and the dominant-active form (Arl2^Q70L^) at E16, three days after IUE, were immunolabelled with tdTomato (Td) and TBR1 (immature neuron marker and labelling CP). (C) Bar graphs (images in A) representing Td+ cell population in the group of control (VZ=13.23 ± 1.85%, SVZ=19.47 ± 3.19%, IZ=45.92 ± 5.44%, CP=21.39 ± 1.58%, n = 4 embryos), human Arl2^WT^ (hArl2^WT^) (VZ=11.13 ± 1.90%, SVZ=18.21 ± 1.98%, IZ=40.25 ± 1.14%, CP=30.41 ± 2.66%, n = 4 embryos), mouse Arl2^WT^ (mArl2^WT^) (VZ=13.92 ± 2.44%, SVZ=20.69 ± 1.70%, IZ=36.09 ± 3.69%, CP=29.30 ± 5.07%, n = 3 embryos) and Arl2^K71R^ mutant (VZ=11.01 ± 3.05%, SVZ=17.18 ± 2.20%, IZ=38.89 ± 1.22%, CP=32.92 ± 6.29%, n = 3 embryos). (D) Bar graphs (images in B) representing Td+ cell population in the group of control (VZ=14.23 ± 2.15%, SVZ=20.15 ± 2.25%, IZ=39.63 ± 3.28%, CP=25.99 ± 4.64%, n = 4 embryos), Arl2^WT^ (VZ=11.13 ± 1.90%, SVZ=18.21 ± 1.98%, IZ=40.25 ± 1.14%, CP=30.41 ± 2.66%, n = 4 embryos), dominant-negative form (Arl2^T30N^) (VZ=15.23 ± 3.51%, SVZ=19.34 ± 2.89%, IZ=65.43 ± 3.78%, CP=0.00 ± 0.00%, n = 4 embryos) and dominant-active form (Arl2^Q70L^) (VZ=15.68 ± 2.44%, SVZ=20.80 ± 1.84%, IZ=63.53 ± 2.86%, CP=0.00 ± 0.00%, n = 4 embryos). (E) Cortical brain sections following overexpression of the dominant-active form (Arl2^Q70L^) at E18, five days after IUE, were immunolabelled with tdTomato (Td) and TBR2 is intermediate progenitor cells marker. TBR1 is immature neuron marker. (F) Quantification graphs representing TBR2+Td+ cells population in the group of control (7.56 ± 0.84%) and Arl2^Q70L^ mutants (42.11 ± 3.45%) in the IZ. (G) Quantification graphs representing TBR1+Td+ cells population in the group of control (8.17 ± 2.59%) and Arl2^Q70L^ mutants (42.91 ± 2.48%) in the IZ. The values represent the mean ± s.d.. Two-Way ANOVA with Multiple comparisons in C and D; Student’s t-test in F and G, n = 3 embryos. Differences were considered significant at *p<0.05, **p<0.01, ***p<0.001 and ****p<0.0001, ns = non-significance. Scale bars; A, B and E = 80 µm, Boxed image for E = 50 µm.

### Overexpression of mutant forms of ARL2 resulted in a defect in neuronal migration

To further elucidate the role of Arl2 in neurogenesis, we tested the effect of overexpression of wildtype Arl2 (Arl2^WT^) as well as two mutant forms of Arl2, namely, the dominant-negative form (Arl2^T30N^) and the dominant-active form (Arl2^Q70L^). Expression of these constructs tagged with Tdtomato (FUtdTW), were driven by the Ubiquitin C (UbC) promotor, in the primary culture of mNPCs *in vitro* by lentivirus (FUtdTW) infection in 48 h culture and pulse-labelled with EdU for 3 h before harvesting the cells. Interestingly, there was a significant increase in the proportion of EdU+ cells in Arl2^WT^ (79.01%) as compared to control (59.06%) (Supplementary Figure 2A and B). In contrast, we observed a significant decrease in the proportion of EdU+ cells in Arl2^T30N^ and Arl2^Q70L^ (36.4% and 27.4%, respectively) as compared to the control group (59.06%) (Supplementary Figure 2A and B). Furthermore, overexpression of Arl2^T30N^ and Arl2^Q70L^ but not Arl2^WT^ caused significant cell death as seen by the increase in caspase-3 staining in mNPCs *in vitro* as compared to control (Supplementary Figure 2A and C; Control = 22.18%; Arl2^WT^ = 30.89%; Arl2^T30N^ = 86.76% and Arl2^Q70L^ = 71.86%). Taken together, our data suggests that Arl2 is required for the proliferation of neural stem cells *in vitro*.

To assess the impact of Arl2 overexpression on the proliferation, differentiation, and migration of mouse neural progenitor cells (mNPCs) during *in vivo* cortical development, we introduced Arl2 and two mutant forms via microinjection into the lateral ventricle of mouse embryos, followed by *in utero* electroporation (IUE) at embryonic day 13 (E13). Remarkably, at E16, overexpression of both mutant forms displayed a significant reduction in Td+ cells migrating to the cortical plate (Figure 2B and D; Arl2^T30N^, VZ = 15.23%, SVZ = 19.34%, IZ = 65.43%, CP = 0.00% and Arl2^Q70L^, VZ = 15.68%, SVZ = 20.80%, IZ = 63.53%, CP = 0.00%) as compared to control (Figure 2B and D; VZ = 14.23%, SVZ = 20.15%, IZ = 39.63%, CP = 25.99%). Our results suggest that overexpression of both Arl2^T30N^ and Arl2^Q70L^ results in a similar phenotype in neuronal migration as Arl2 KD.

Surprisingly, we find that overexpression of Arl2^T30N^ and Arl2^Q70L^ but not Arl2^WT^ at E15, two days after IUE, showed a significant increase in the proportion of phospho-histone H3-positive (PH3+) cells in the VZ as compared to control (Supplementary Figure 2D-E; control 3.31%, Arl2^WT^ 2.94%, Arl2^T30N^ 8.33%, Arl2^Q70L^ 9.43%). This increase in PH3+ cells may be due to mitotic defects or over proliferation of radial glial cells. To distinguish these two possibilities, we examined the spindle poles marked by gamma-tubulin in these cells to assess their cell divisions. Indeed, overexpression of Arl2^T30N^ and Arl2^Q70L^ caused a significant increase in mitotic defects (cell cycle arrest), with defective spindle formation as compared to control and Arl2^WT^ (Supplementary Figure 3A). Furthermore, similar to our *in vitro* results, there was a significant increase in the proportion of caspase-3+ cells in the IZ in Arl2^Q70L^ (38.52%) as compared to control (5.92%; Supplementary Figure 3B and C). These data suggest that the migration and proliferation defects observed in Arl2 mutants are possibly due to mitotic defects, eventually leading to cell death.

To further test whether Arl2 affects neuronal migration, we examined TBR2, a transcription factor that marks the transition from radial glial cells to intermediate progenitor cells (Figure 2E-G). At E18, 5 days after IUE, we find that majority of the TBR2+Td+ cells have already migrated to the CP in control mouse brains, with few TBR2+Td+ cells still present in the IZ (Figure 2E-G). Interestingly, in Arl2^Q70L^ mutants a large population of TBR2+Td+ cells still remained in the IZ (Figure 2F, 42.11 ± 3.45%) with few to no cells migrating to the CP as compared to control (7.56 ± 0.84%). Similarly, a vast majority of neuronal (TBR1+) were still retained in the IZ in Arl2^Q70L^ mutant mouse brains (Figure 2G; 42.91 ± 2.48%) as compared to control (8.17 ± 2.59%). In contrast, the expression of NeuroD2, a neuronal marker found in immature neurons, is significantly increased in Arl2^WT^ (30.77 ± 2.93%) but dramatically reduced in Arl2^Q70L^ (6.75 ± 2.69%) 3 days after IUE as compared to control (20.57 ± 1.36%) (Supplementary Figure 3D-E).

Taken together, Arl2 dysfunction resulted in a defective migration of neuronal cells.

### Loss of Arl2 results in a significant reduction of centrosomal microtubule growth in mNPCs *in vitro*

Arl2 regulates centrosomal microtubule nucleation and growth in various cell types including *Drosophila* NSCs [21, 22, 24]. We sought to investigate whether mouse Arl2 plays a conserved role in microtubule growth in mNPCs. To this end we performed microtubule regrowth assay, where in, mNPCs were synchronized in the S phase using thymidine and microtubules were depolymerized by nocodazole treatment in these cells. Microtubule regrowth labelled by α-tubulin was assessed in a time-course experiment following wash out of nocodazole (Figure 3A). Before nocodazole treatment, shArl2 cells showed a slight but not significant reduction of microtubule intensity (97.9 A.U.) as compared to control (Figure 3B-C, 122.70 A.U.). Microtubules were efficiently depolymerized in both control and Arl2 KD cells (t = 0), as only weak residual microtubules labelled by α-tubulin were seen at the centrosome following nocodazole treatment (Figure 3B-C, shCtrl: 30.21 A.U. vs shArl2: 31.63 A.U.). In control, robust microtubules were observed around the centrosome, at various time points following recovery (Fig 3B-C). In contrast, Arl2 KD in mNPCs reassembled less microtubule mass after 5 mins recovery (Fig. 3B-C; shCtrl: 68.27 A.U. vs shArl2: 39.47 A.U.), and after 10 min (Fig. 3B-C; shCtrl: 95.96 A.U. vs shArl2: 67.50 A.U.). Interestingly, even after 30 minutes of recovery, Arl2 KD cells were still unable to recover their microtubule mass as compared to control (Fig. 3B-C; shCtrl: 110.87 A.U. vs shArl2: 71.63 A.U.). These results suggest that Arl2 promotes microtubule growth in mNPCs.

**Fig. 3.**
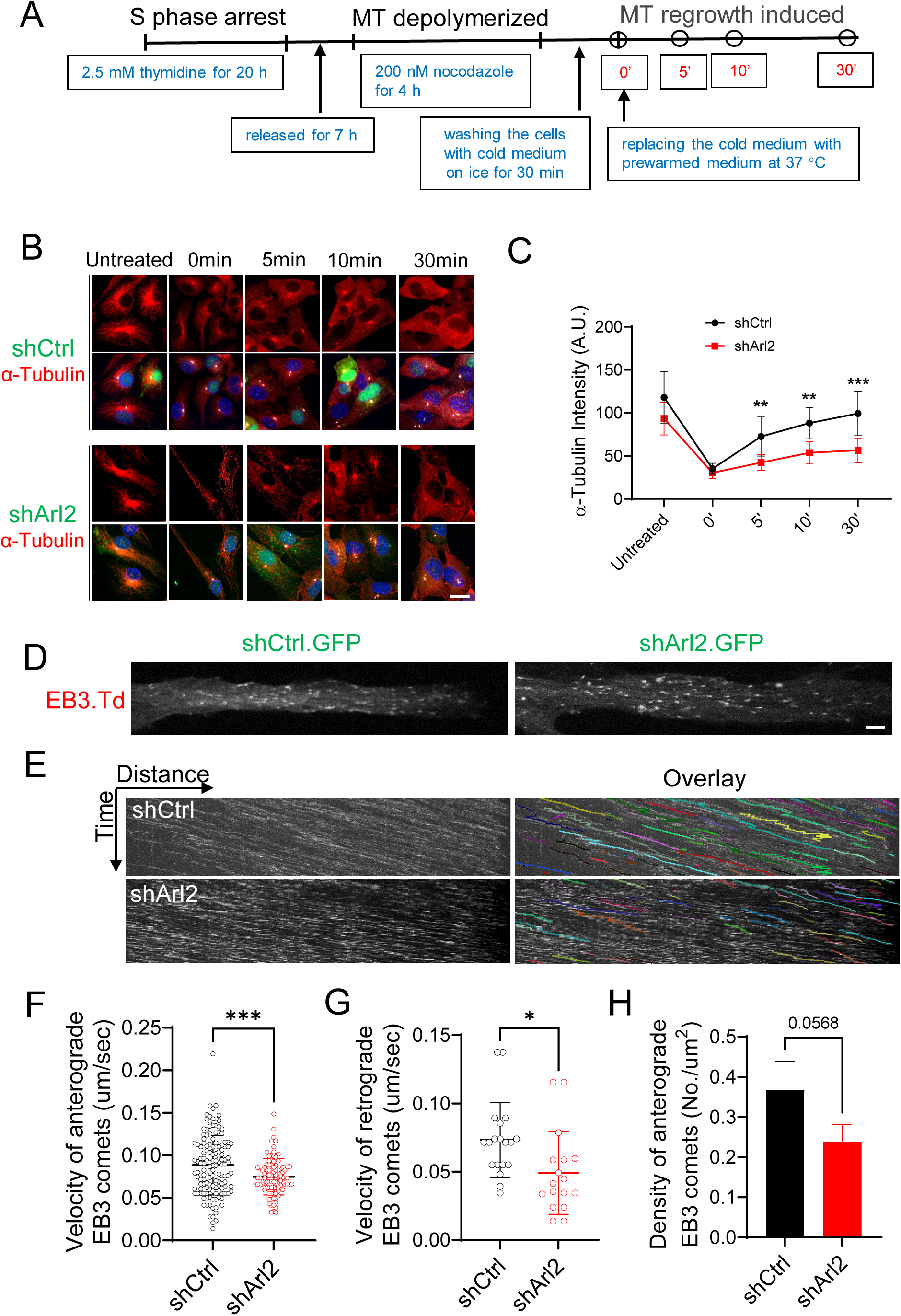
Loss of Arl2 results in a significant reduction of centrosomal microtubule growth in mNPCs *in vitro*. (A) Schematic representation of centrosomal microtubule regrowth assay. (B) Immunostaining micrographs showing the microtubule regrowth labelled by α-tubulin within the time course (0, 5, 10, and 30 min) in both shCtrl and shArl2 groups. (C) Line graph representing α-tubulin intensity in shArl2 group (Untreated =97.9 ± 26.30, 0 min = 31.63 ± 7.26, 5 min = 39.47 ± 9.78, 10 min = 67.50 ± 12.87, and 30 min =71.63 ± 12.48) compared to the control (Untreated =122.70 ± 27.32, 0 min = 30.21 ± 9.20, 5 min = 68.27 ± 18.19, 10 min = 95.96 ± 18.40, and 30 min =110.87 ± 18.38), (Unit = A.U.). The values represent the mean ± s.d.. Multiple t-test in C, n = 3. Differences were considered significant at **p<0.01, ***p<0.001. (D) Live imaging micrograph to track the growing ends of microtubules by using the plus-end microtubule binding protein EB3 tagged with Tdtomato (Td) in mNPCs in both shCtrl and shArl2 groups. (E) Kymographs showing the EB3-Td comets movement in mNPCs in both shCtrl and shArl2 groups. (F, G & H) Quantification graphs representing the velocity of anterograde EB3 comets (shCtrl: 0.09 ± 0.03 µm/sec vs shArl2: 0.07 ± 0.02 µm/sec), the velocity of retrograde EB3 comets (shCtrl: 0.07 ± 0.03 µm/sec vs shArl2: 0.05 ± 0.03 µm/sec) and the total density of EB3 comets (shCtrl: 0.30 ± 0.05 No./µm2 vs shArl2: 0.24 ± 0.04 No./µm2). The values represent the mean ± s.d.. Student’s t-test in F, G and H, n = 3. Differences were considered significant at *p<0.05, ***p<0.001. Scale bars; B = 10 µm, D = 1 µm.

To further determine the role of Arl2 in microtubule growth of mNPCs, we performed live imaging to track the growing ends of microtubules by using the plus-end microtubule binding protein EB3, which is enriched in the nervous system [31]. Remarkably, knock down Arl2 (shArl2 marked by GFP) in mNPCs expressing EB3-tdTomato resulted in a significant reduction in velocity of both anterograde (0.07 µm/sec) and retrograde EB3 comets (0.05 µm/sec) as compared to control (Figure 3D-G; Movie S3; anterograde, 0.09 µm/sec; retrograde, 0.07 µm/sec). Furthermore, the total density of EB3 comets was notably reduced as compared to the control (Figure 3D, E and H; Movie S3; shCtrl = 0.30 No./µm^2^; shArl2 = 0.24 No./µm^2^). Taken together, our data suggests that Arl2 is critical for microtubule growth in mNPCs.

### Loss of Arl2 led to a shift of symmetric division to asymmetric division of mNPCs and alters the mNPC differentiation

Next, we investigated whether the defects in mNPC proliferation and differentiation upon Arl2 KD was caused by an imbalance between the symmetric and asymmetric division of NPCs. Control and Arl2 shRNA were introduced via microinjection into the lateral ventricle of mouse embryos, followed by IUE at embryonic day 13 (E13). At E14, one day after IUE, wild-type radial glial cells in the VZ can undergo both symmetric and asymmetric divisions, depending on the plane of division (Figure 4A-C). Interestingly, loss of Arl2 causes a significantly larger population of radial glial cells to divide with an oblique orientation (30°-60°) at the cleavage furrow (31.36%) and fewer radial glial cells (46.94%) divide in the vertical division plane (60°-90°) as compared to control (30°-60° = 21.08% and 60°-90° = 56.15%; Figure 4A-C). There was no obvious change the population of radial glial cells that divide in the horizontal division plane (0°-30°; shCtrl = 22.78% vs shArl2 = 21.70%). Thus, Arl2-depleted NPCs divide asymmetrically more frequently than symmetrically as compared to control, which may account for the defects in NPC proliferation and differentiation observed upon loss of Arl2. This data suggests that Arl2 regulates radial glia proliferation possibly by regulating proper centrosomal localization, mitotic spindle formation and cell cycle progression.

**Fig. 4.**
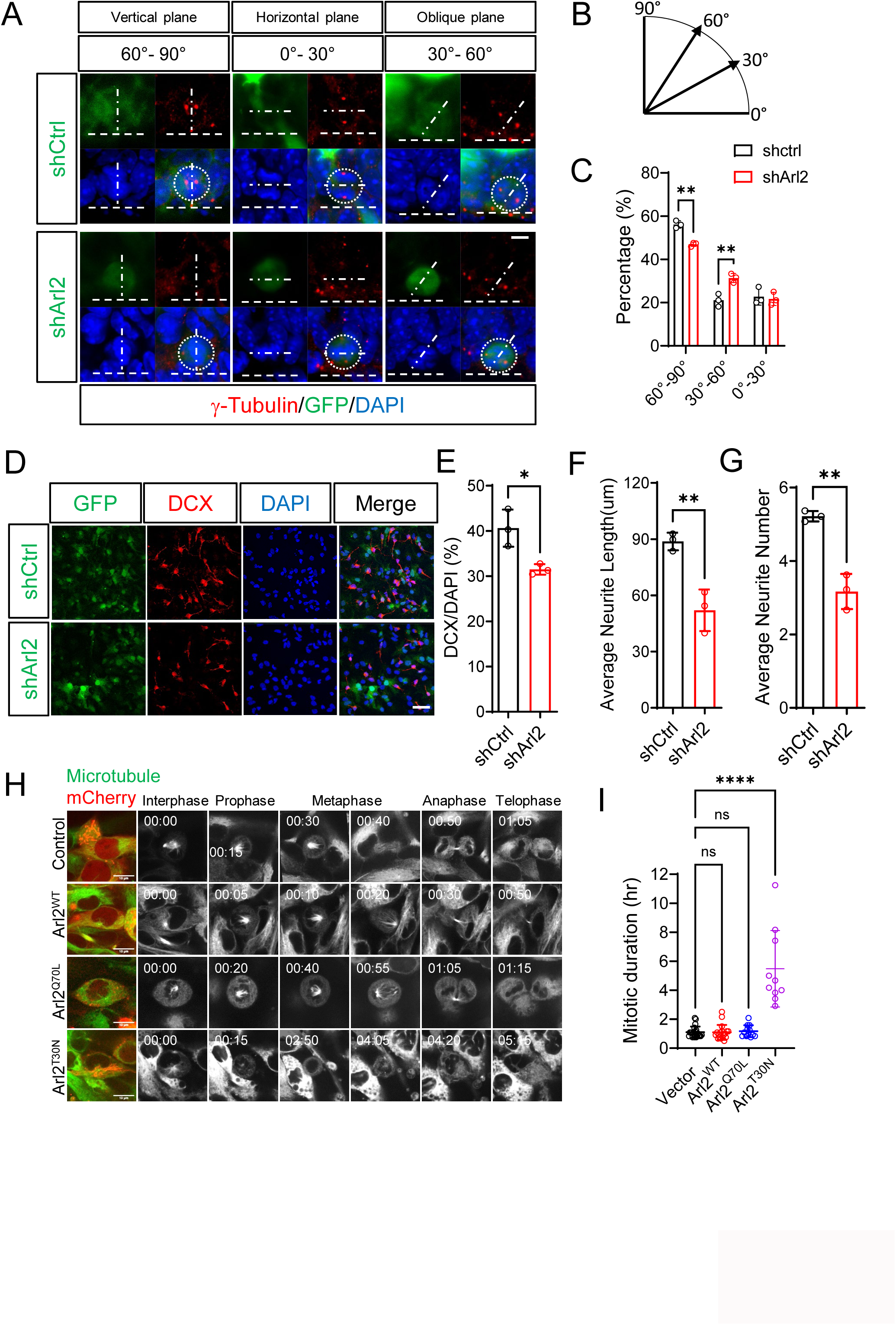
Loss of Arl2 led to a shift in symmetric division to asymmetric division of mNPCs and alters the mNPC differentiation. (A & B) Immunostaining micrographs showing transition from symmetric to asymmetric cell division (brain section) after 2 days IUE. The centrosome of mitotic cells was labelled using antibodies against γ-Tubulin which organize the mitotic spindle. The orientation of the mitotic centrosome at the cleavage furrow relative to the brain ventricular surface (horizontal dash line) was determined. The symmetric division was determined by the vertical cleavage plain (60-90°), and the asymmetric division was determined by the horizontal cleavage plain (0-30°) and oblique cleavage plain (30-60°). (C) Bar graph representing the population of radial glial cells to divide with horizontal cleavage plain (0-30° =21.70 ± 3.03%), oblique cleavage plain (30°-60° = 31.36 ± 2.09%) and vertical cleavage plain (60°-90° = 46.94 ± 0.97%) in shArl2 group as compared to control (0°-30° = 22.78 ± 3.93%, 30°-60° = 21.08 ± 2.91% and 60°-90° = 56.15 ± 1.68%). The values represent the mean ± s.d.. Multiple t-test in C, n = 3. Differences were considered significant at **p<0.01. (D) Immunostaining micrographs showing the DCX+ immature neurons after 5 days mNPC differentiation assay in both shCtrl and shArl2 groups. (E, F &G) Bar graphs representing the population of DCX-positive cells (shCtrl: 40.63 ± 4.11%, shArl2: 31.48 ± 1.72%); the average neurite length (shCtrl: 88.88 ± 4.69 µm, shArl2: 53.60 ± 11.91 µm) and the average neurite number (shCtrl: 5.22 ± 0.14, shArl2: 3.27 ± 0.52). The values represent the mean ± s.d.. Student’s t-test in F, G and H, n = 3. Differences were considered significant at *p<0.05, **p<0.01. (H) Time series of mNPCs *in vitro* using Viafluor-488 live cell microtubule staining kit (Biotium, #70062) in Arl2^WT^, Arl2^Q70L^ and Arl2^T30N^. (I) Quantification graph showing the average time taken for a single mitotic cycle in control = 1.10 ± 0.39 hours, Arl2^WT^ = 1.09 ± 0.51 hours, Arl2^Q70L^ = 1.17 ± 0.41 hours, Arl2^T30N^ = 5.48 ± 2.64 hours in mNPCs overexpressing Arl2^WT^, Arl2^Q70L^ and Arl2^T30N^. Scale bars; A = 5 µm; D = 50 µm; H = 10 µm.

To investigate whether Arl2 regulates neuronal differentiation of mNPCs, we examined Doublecortin (DCX), a neuronal marker that is expressed in neuronal precursor cells and immature neurons. Interestingly, Arl2 KD in mNPCs led to a significant reduction in the population of DCX-positive cells (31.48%) as compare to control (40.63%; Figure 4D-E). Furthermore, the average neurite length (53.60 µm) as well as average neurite number (3.27) were significantly reduced in Arl2 KD as compared to control (neurite length = 88.88 µm, neurite number = 5.22; Figure 4D, F-G), suggesting that loss of Arl2 affects neuronal differentiation of mNPCs.

To further understand how Arl2 affects mNPC proliferation, we performed live imaging of mNPCs *in vitro* using the Viafluor-488 live cell microtubule staining kit (Biotium, #70062). In control mNPCs the average time taken for a single mitotic cycle was 1.10 hours (Figure 4H-I; Movie S4). Interestingly, mNPCs expressing Arl2^T30N^ but not Arl2^Q70L^ or Arl2-WT caused a significant increase in the mitotic duration as compared to the control (Figure 4H-I; Movie S4; average time, Arl2^T30N^ = 5.48, Arl2^Q70L^ = 1.17, Arl2^WT^ = 1.09 hours), suggesting an additional defect in the cell cycle progression caused by this mutant form.

### Arl2 regulates the neuritogenesis of mouse primary neurons

To determine whether neuronal migration defects caused by Arl2 dysfunction is due to impaired neuritogenesis, we examined the effect of Arl2^T30N^ and Arl2^Q71L^ in mouse primary neurons *in vitro*. Interestingly, overexpression of Arl2^T30N^ and Arl2^Q71L^ dramatically affected neuronal morphology, as fewer and shorter neurites were observed in these neurons as compared to control (Supplementary Figure 4A). The total intersection number as measured by Scholl’s analysis was significantly reduced in Arl2^Q71L^ (11 ± 1.80) and Arl2^T30N^ (8.83 ± 1.04) as compared to control (38 ± 4.58; Supplementary Figure 4A-C). These data suggest that neuronal migration defects caused by Arl2 dysfunction is possibly due to its role in regulating neurite outgrowth.

### Mitochondria defect is not the major cause of neurogenesis deficit observed in Arl2 dysfunction

Given the importance of Arl2 in regulating both microtubule growth and mitochondrial functions [21, 30] we sought to pinpoint the mechanism underlying the role of Arl2 during neurogenesis. To this end, we generated the Lys71 to Arg71 (Arl2^K71R^) mutation in mouse Arl2 which is known to cause mitochondrial fragmentation and immobility without disruption in microtubule assembly [32]. As expected, Arl2^K71R^ overexpression showed fragmented mitochondria with shortened mitochondrial length (2.41 ± 0.99 µm) as compared to control (5.53 ± 3.78 µm; Supplementary Figure 4D-E) in mouse NPCs *in vitro*. Interestingly, at day 3 after IUE during *in vivo* cortical development, overexpression of mouse Arl2^K71R^ mutant, resulted in a significant increase in the number of Td+ cells migrating to the CP (VZ = 11.01%, SVZ = 17.18%, IZ = 38.89%, CP = 32.92%) as compared to control (VZ = 13.23%, SVZ = 19.47%, IZ = 45.92%, CP = 21.39%; Figure 2A and C), which mimicked the effect of overexpression of the wild-type Arl2 on cortical development (Figure 2).

Consistent with previous reported role of Arl2 in mitochondria fusion [26], Arl2^Q70L^ overexpression in mNPCs caused a dramatic decrease in the number of cells with tubular mitochondria, indicating an increase in mitochondria fusion, while Arl2^T30N^ overexpression resulted in a severe mitochondrial fragmentation as compared to the control or Arl2^WT^ overexpression (Supplementary Figure 5A and B; Movie S5). Furthermore, neither shArl2 KD nor overexpression of Arl2, showed any change in mitochondrial morphology in mouse brains 4 days after IUE as compared to control (Supplementary Figure 5C). Taken together, mitochondrial defects were not the primary cause for migration deficits observed in Arl2 dysfunction.

### Overexpression of Arl2 mutant forms leads to defects in microtubule growth in mNPCs

#### in vitro

Next, we performed microtubule regrowth assay for mNPCs overexpressing Arl2^WT^, Arl2^Q70L^ and Arl2^T30N^. Both control and Arl2^WT^ cells treated with nocodazole (t = 0s) showed weak residual microtubules at the centrosome, suggesting an efficient microtubule depolymerization (Figure 5A-B, control: 30.21 ± 9.20; Arl2^WT^: 47.30 ± 8.64). Remarkably, after 5 minutes of recovery Arl2^WT^ mNPCs displayed more abundant microtubule density than control (Figure 5A-B, control: 61.40 ± 18.00 A.U.; Arl2^WT^: 76.31 ± 24.34 A.U.). Even after 30 minutes of recovery, Arl2^WT^ mNPCs still showed significantly higher microtubule density as compared to control (Figure 5A-B, control: 96.65 ± 19.09 A.U.; Arl2^WT^: 110.74 ± 17.96 A.U.), suggesting that overexpression of Arl2^WT^ likely leads to overgrowth of microtubules in mNPCs. Similarly, overexpression of Arl2^K71R^ caused a significant increase in overall microtubule density, even after 30 minutes of recovery (0 min = 62.56 ± 15.69 A.U., 5 min = 90.77 ± 14.18 A.U., 10 min = 120.69 ± 19.26 A.U., and 30 min =115.93 ± 20.06 A.U.), as compared to control recovery (33.82 ± 10.65 A.U., 5 min = 63.64 ± 15.80 A.U., 10 min = 79.52 ± 14.88 A.U., and 30 min = 81.37 ± 14.90 A.U.; Supplementary Figure 4F), suggesting that Arl2^K71R^ behaves as the Arl2 wild-type form. In contrast, overexpression of Arl2^Q70L^ and Arl2^T30N^ in mNPCs reassembled significantly lesser microtubule mass, at various time points following recovery as compared to control (Fig 5A-B; Arl2^T30N^, Untreated =73.65 ± 7.14 A.U., 0 min = 40.34 ± 5.38 A.U., 5 min = 43.97 ± 13.29 A.U., 10 min = 31.33 ± 9.64 A.U., and 30 min =40.75 ± 13.64 A.U.; Arl2^Q70L^, Untreated =79.76 ± 13.13 A.U., 0 min = 26.45 ± 10.44 A.U., 5 min = 52.94 ± 21.58 A.U., 10 min = 43.20 ± 7.83 A.U., and 30 min =65.82 ± 14.84 A.U.). These results further support that Arl2 promotes microtubule growth in mNPCs.

**Fig. 5.**
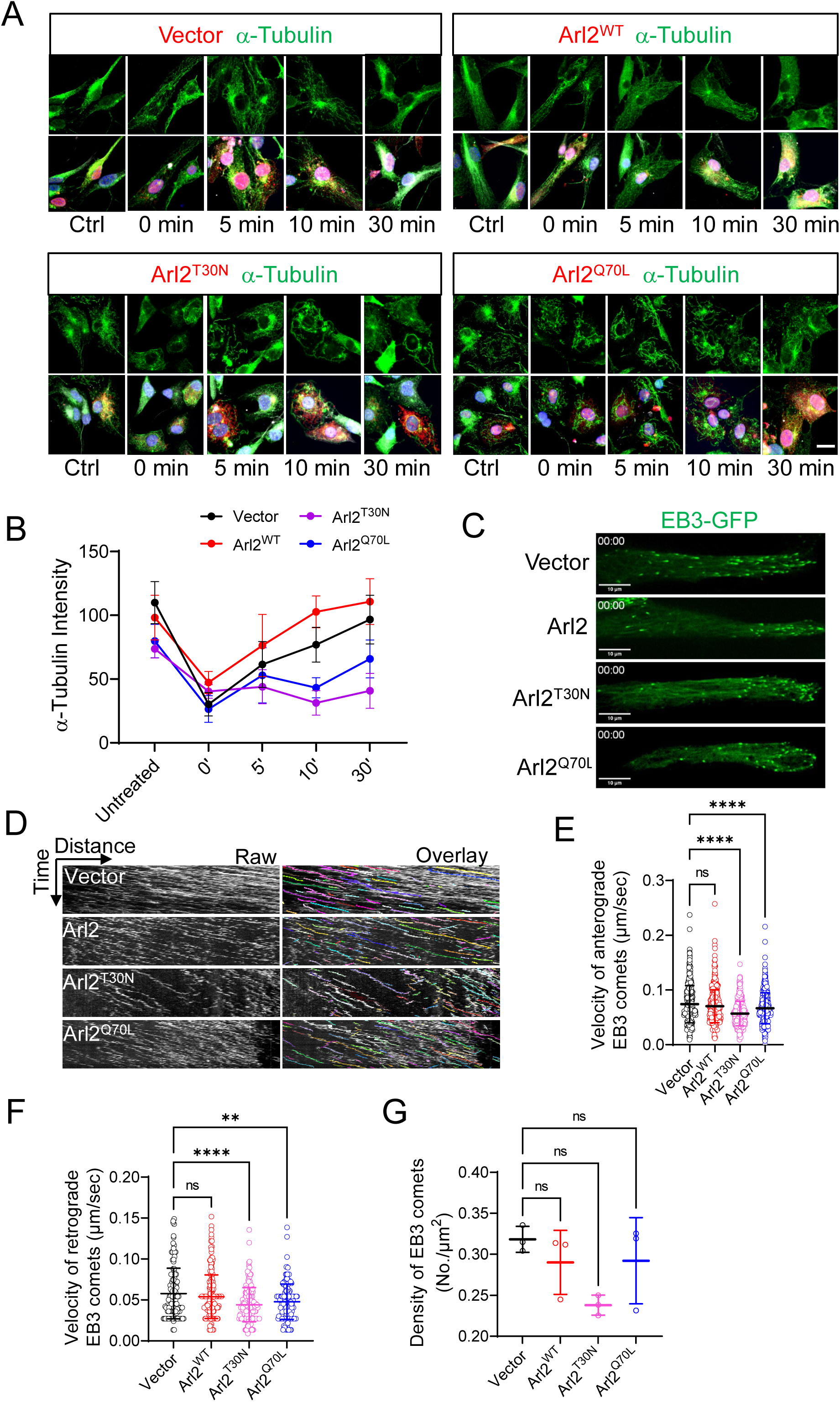
Overexpression of mutant forms of Arl2 leads to defects in microtubule growth in mNPCs. (A) Immunostaining micrographs showing microtubule regrowth labelled by α-tubulin within the time course (0, 5, 10, and 30 min) in Arl2^WT^, Arl2^T30N^ and Arl2^Q70L^ groups in mNPCs. (B) Line graph representing α-tubulin intensity in Arl2^WT^ group (Untreated =98.22 ± 17.51, 0 min = 47.30 ± 8.64, 5 min = 76.31 ± 24.34, 10 min = 102.69 ± 12.52, and 30 min =110.74 ± 17.96), Arl2^T30N^ group (Untreated =73.65 ± 7.14, 0 min = 40.34 ± 5.38, 5 min = 43.97 ± 13.29, 10 min = 31.33 ± 9.64, and 30 min =40.75 ± 13.64), Arl2^Q70L^ group (Untreated =79.76 ± 13.13, 0 min = 26.45 ± 10.44, 5 min = 52.94 ± 21.58, 10 min = 43.20 ± 7.83, and 30 min =65.82 ± 14.84) compared to the control (Untreated =109.90 ± 16.48, 0 min = 30.21 ± 9.20, 5 min = 61.40 ± 18.00, 10 min = 76.94 ± 13.60, and 30 min =96.65 ± 19.09) in mNPCs. The values represent the mean ± s.d., Unit = A.U.. Two-Way ANOVA with Multiple comparisons in B, n = 3. Differences were considered significant at *p<0.05, **p<0.01, ***p<0.001 and ****p<0.0001, ns = non-significance. (C) Live imaging micrograph to track the growing ends of microtubules by using the plus-end microtubule binding protein EB3 tagged with GFP in mNPCs in Arl2^WT^, Arl2^T30N^ and Arl2^Q70L^ groups. (D) Kymographs showing the EB3-GFP comets movement in mNPCs in Arl2^WT^, Arl2^T30N^ and Arl2^Q70L^ groups. (E, F & G) Quantification graphs representing the velocity of anterograde EB3 comets (Control: 0.074 ± 0.034 µm/sec, Arl2^WT^: 0.070 ± 0.03 µm/sec; Arl2^T30N^: 0.057 ± 0.022 µm/sec and Arl2^Q70L^: 0.067 ± 0.028 µm/sec), the velocity of retrograde EB3 comets (Control: 0.058 ± 0.031 µm/sec, Arl2^WT^: 0.054 ± 0.027 µm/sec; Arl2^T30N^: 0.044 ± 0.021 µm/sec and Arl2^Q70L^: 0.048 ± 0.021 µm/sec) and the total density of EB3 comets (Control: 0.32 ± 0.02 No./µm^2^, Arl2^WT^: 0.29 ± 0.04 No./µm^2^; Arl2^T30N^: 0.24 ± 0.01 No./µm^2^ and Arl2^Q70L^: 0.29 ± 0.05 No./µm^2^, n = 3). The values represent the mean ± s.d.. One-Way ANOVA in E, F and G. Differences were considered significant at **p<0.01 and ****p<0.0001, ns = non-significance. Scale bars; A = 5 µm; C = 10 µm.

To further analyse the effect of Arl2 on microtubule growth, we performed live imaging in mNPCs overexpressing Arl2^WT^, Arl2^Q70L^ or Arl2^T30N^. Remarkably, overexpression of Arl2^WT^ showed a significant increase in the intensity of microtubules as compared to control (Supplementary Figure 5A). In contrast, both Arl2^Q70L^ and Arl2^T30N^ showed a significant reduction in the intensity of microtubules as compared to control (Supplementary Figure 5A).

To further validate the role of Arl2 in microtubule growth, we performed live imaging of EB3-GFP comets in Arl2^WT^, Arl2^Q70L^ and Arl2^T30N^ mNPCs. Similar to Arl2 KD, overexpression of Arl2^Q70L^ and Arl2^T30N^ resulted in a significant reduction in velocity in anterograde (Arl2^T30N^, 0.057 ± 0.022 µm/sec; Arl2^Q70L^, 0.067 ± 0.028 µm/sec) and retrograde EB3 comets (Arl2^T30N^, 0.044 ± 0.021 µm/sec and Arl2^Q70L^, 0.048 ± 0.021 µm/sec) as compared to control (Figure 5C-F; Movie S6; anterograde, (0.074 ± 0.034 µm/sec), retrograde, (0.058 ± 0.031 µm/sec). The total density of EB3 comets appeared to be normal by overexpression of Arl2^WT^, Arl2^Q70L^ or Arl2^T30N^ (Figure 5C-D, G; Movie S6). Furthermore, overexpression of Arl2^WT^ showed no obvious change in movements of EB3-GFP comets (Figure 5D-G), likely due to its weaker effects than overexpression Arl2 mutant forms. Taken together, these observations indicate that Arl2 regulates microtubule growth in mNPCs.

### Arl2 localizes to the PCM of the centrosomes and facilitates γ-tubulin localization at the centrosomes in mNPCs

Arl2 localizes to the centrosomes in different cell types including HEK and CHO cells and presumably localizes to PCM [20]. To determine whether Arl2 localizes to the PCM in HEK293 cells, we examined the ultrastructure of Arl2 (Arl2-HA) and Cdk5rap2 (Cdk5rap2-Myc), a centrosomal protein which is involved in microtubule organization [33], using super resolution microscopy (Figure 6A). Remarkably, Arl2 and Cdk5rap2 formed ring-like structures that co-localized with one another at the centrosome co-labelled by γ-tubulin in interphase and metaphase cells (Figure 6A). This observation suggests that Arl2 indeed localizes to the PCM. Furthermore, knocking down of Arl2 resulted in a significant decrease in γ-tubulin intensity at the centrosomes in metaphase of mNPCs (Figure 6B, D; 68.59 ± 9.31 A.U.) and interphase of mNPCs (supplementary Figure 6C, E; 52.83 ± 10.22 A.U.) as compared to control (metaphase: 114.9 ± 24.88 A.U., interphase: 80.47 ± 23.81 A.U., respectively). These data suggest that Arl2 is a centrosomal protein required for centrosomal assembly in mNPCs.

**Fig. 6.**
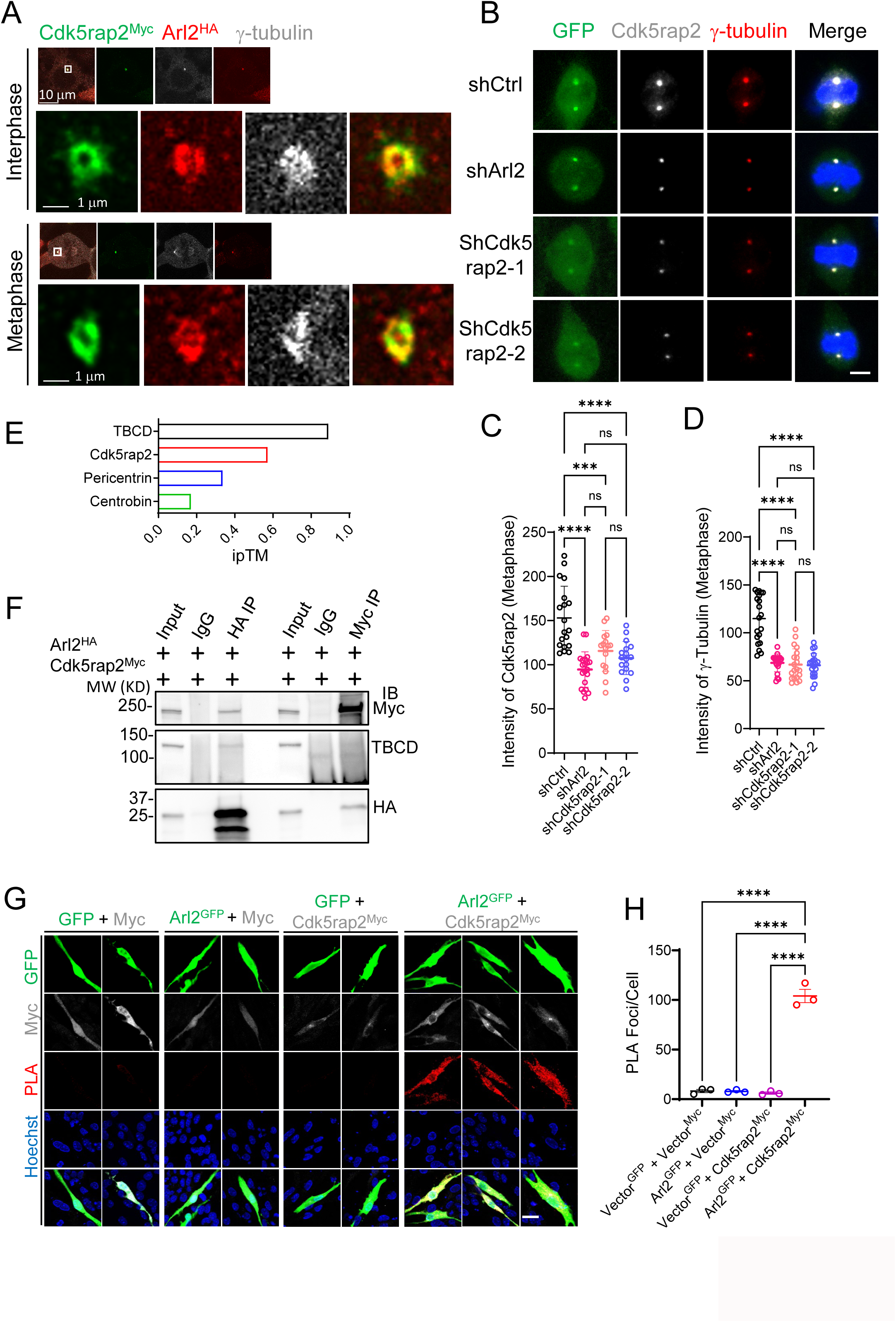
Arl2 localizes to the centrosomes and is required for γ-tubulin localization at the centrosomes in mNPCs. (A) Immunostaining micrographs of HEK293 cells co-transfected with Arl2-HA and Cdk5rap2-Myc were imaged using super resolution microscopy labelled for γ-tubulin, Myc, and HA. (B) Immunostaining micrographs showing Cdk5rap2 and γ-tubulin in shCtrl, shArl2 and shCdk5rap2 groups in mNPCs. (C) Quantification graph showing Cdk5rap2 intensity at metaphase of mNPCs (94.5 ± 20.06, n = 3 batches with 20 cells) upon Arl2 knockdown and (shCdk5rap2-1 = 115.6 ± 23.13, n = 3 batches with 16 cells; shCdk5rap2-2 = 107.5 ± 18.77, n = 3 batches with 17 cells) upon Cdk5rap2 knockdown as compared to shCtrl (153.0 ± 35.96, n = 3 batches with 19 cells). (D) Quantification graph showing γ-tubulin intensity at metaphase of mNPCs (68.59 ± 9.31, n = 3 batches with 20 cells) upon Arl2 knockdown and (shCdk5rap2-1 = 66.85 ± 16.44, n = 3 batches with 20 cells; shCdk5rap2-2 = 66.11 ± 12.33, n = 3 batches with 20 cells) upon Cdk5rap2 knockdown as compared to shCtrl (114.9 ± 24.88, n = 3 batches with 20 cells). The values represent the mean ± s.d.. One-Way ANOVA in C and D. Differences were considered significant at ***p<0.001 and ****p<0.0001, ns = non-significance. (E) Bar graph showing Alpha fold multimer interaction prediction of Arl2 and Cdk5rap2, Arl2 and Pericentrin (PCNT), Arl2 and Centrobin with an ipTM score of 0.57, 0.34, 0.17, respectively, compared with Tubulin folding cofactor D (TBCD), a known interactor of Arl2 with an ipTM score of 0.89. (F) Co-immunoprecipitation by over-expressing Arl2 (HA-Arl2) and Cdk5rap2 (Myc-Cdk5rap2) in HEK293 cells. Following precipitation with a HA antibody, the resulting protein complexes exhibited an anticipated 37 kD band corresponding to HA-Arl2 as well as 250 kD band corresponding to Myc-Cdk5rap2. TBCD was used as positive control which also co-immunoprecipitated following precipitation with a HA antibody. Similarly, following precipitation with Myc antibody, bands corresponding Myc-Cdk5rap2 and HA-Arl2 were observed. (G) Proximity Ligation Assay (PLA) showing over-expressing Arl2 (Arl2-GFP) and Cdk5rap2 (Myc-Cdk5rap2) (Vector-GFP and Myc-Vector, Arl2-GFP and Myc-Vector, Vector-GFP and Myc-Cdk5rap2, Arl2-GFP and Myc-Cdk5rap2) in mNPCs. (H) Quantification graph of the PLA foci per cell with no red dot, weak red dots, and strong red dots for (G). Vector-GFP and Myc-Vector, 8.17 ± 2.75; Arl2-GFP and Myc-Vector, 7.83 ± 1.44; Vector-GFP and Myc-Cdk5rap2, 6.11 ± 1.99; Arl2-GFP and Myc-Cdk5rap2, 104.00 ± 11.53; n = 3). The values represent the mean ± s.d.. One-Way ANOVA in H. Differences were considered significant at ****p<0.0001. Scale bars; A = 10 µm; Boxed image for A = 1 µm; B = 5 µm; G = 40 µm.

### Arl2 interacts with the centrosomal protein Cdk5rap2

Next, we explored whether Arl2 and Cdk5rap2 can interact with each other. Alpha-fold multimer is emerging as a powerful and accurate approach for *in silico* prediction of protein-protein interactions based on deep learning method [34-36]. Using Alpha-fold multimer, Tubulin folding cofactor D (TBCD), a known strong interactor of Arl2, had an ipTM score of 0.89, indicating the reliability of this approach. We found that Cdk5rap2 had an ipTM score of 0.57 suggesting that Cdk5rap2 is a strong candidate of Arl2-interacting protein (Figure 6E).

Pericentrin, another centrosomal protein required for cortical development, also interacts with and recruits Cdk5rap2 to the centrosome in the mNPCs [16]. Interestingly, Alpha-fold multimer also predicts that Arl2 can potentially interact with Pericentrin with an ipTM score of 0.33 (Figure 6E). Centrobin, a centriolar protein as a negative control for the interaction testing was predicted not to interact with Cdk5rap2 with an ipTM score of 0.17 by Alpha-fold multimer (Figure 6E).

To validate the predicted interaction between Arl2 and Cdk5rap2, we performed co-immunoprecipitation by over-expressing Arl2 (HA-Arl2) and Cdk5rap2 (Myc-Cdk5rap2) in HEK293 cells (Figure 6F). Following immunoprecipitation with a HA antibody, the resulting protein complexes exhibited an anticipated 37 kD band corresponding to HA-Arl2 as well as 250 kD band corresponding to Myc-Cdk5rap2, suggesting that Arl2 and Cdk5rap2 physically associate with each other (Figure 6F). TBCD was used as positive control which also co-immunoprecipitated following precipitation with a HA antibody (Figure 6F). Similarly, following precipitation with Myc antibody, bands corresponding Myc-Cdk5rap2 and HA-Arl2 were observed further confirming the interaction between Cdk5rap2 and Arl2 (Figure 6F).

To further validate the association between Arl2 and Cdk5rap2, we employed proximity ligation assay (PLA), a technique that enables the detection of protein-protein interactions with high specificity and sensitivity [37]. We co-expressed various proteins tagged with Myc or GFP in mNPCs and quantified PLA foci that indicated protein-protein interactions (Figure 6G-H). The vast majority of mNPCs co-expressing both Myc or GFP controls displayed weak fluorescence signal of merely a few PLA foci (Figure 6G-H; 8.17 ± 2.75). Similarly, the vast majority of cells co-expressing Arl2-GFP with control Myc or Myc-Cdk5rap2 with control GFP displayed few PLA puncta per cell 7.83 ± 1.44; Vector-GFP and Myc-Cdk5rap2, 6.11 ± 1.99), under each co-expression condition, respectively (Figure 6G-H). By contrast, mNPCs co-expressing Arl2-GFP and Myc-Cdk5rap2 displayed strong signal with a plethora of PLA foci (Figure 6G-H; 104 ± 11.5).

Taken together, our data indicates that Arl2 and Cdk5rap2 can physically interact with each other.

### Cdk5rap2 affects neuronal migration and proliferation *in vitro* and *in vivo*, similar to Arl2 loss-of-function

Cdk5rap2 maintains NPC pool in the developing Neocortex [16]. We found similar neurogenesis defects following knocking down of Cdk5rap2 (Supplementary Figure 6).

Upon Cdk5rap2 KD in mNPCs by two independent shCdk5rap2-1 and shCdk5rap2-2 tagged with GFP, Cdk5rap2 protein level detected by anti-Cdk5rap2 antibodies in Western blot (WB) was reduced to 26% in shCdk5rap2-1 and 32% in shCdk5rap2-2, respectively, compared with the control (Supplementary Figure 6A-B), suggesting efficient knockdowns by both shCdk5rap2. Silencing endogenous Cdk5rap2 expression in the primary culture of mNPCs *in vitro* by lentivirus (pPurGreen) infection in 48 h culture resulted in a significant decrease in the intensity of Cdk5rap2 at the centrosomes in metaphase mNPCs (shCdk5rap2-1 = 115.6 A.U.; shCdk5rap2-2 = 107.5 A.U., respectively) as compared to control (Figure 6B, C; metaphase: 153.0 A.U.). Similarly, the intensity of Cdk5rap2 at the centrosomes in interphase mNPCs (shCdk5rap2-1 = 34.73 A.U.; shCdk5rap2-2 = 44.53 A.U., respectively) was significantly reduced as compared to control (Supplementary Figure 6B, C; interphase: 77.37 A.U.). Furthermore, there was a significant reduction in γ-tubulin intensity upon Cdk5rap2 KD (shCdk5rap2-1 = 66.85 A.U.; shCdk5rap2-2 = 66.11 A.U., respectively) in metaphase mNPCs as well as at the centrosomes in interphase mNPCs (shCdk5rap2-1 = 32.02 A.U.; shCdk5rap2-2 = 36.29 A.U., respectively) at the as compared to control (metaphase: 114.9 A.U.; interphase: 80.47 A.U.; Figure 6B, C and Supplementary Figure 6B, C). Remarkably, the proportion of EdU+ cells in the primary culture of mNPCs was dramatically reduced upon Cdk5rap2 KD (shCdk5rap2-1 = 26.42 ± 7.89%; shCdk5rap2-2 = 34.58 ± 5.72%, respectively) as compared to the control group (shCtrl = 58.84 ± 6.06%) (Supplementary Figure 6F, G).

We introduced Cdk5rap2 shRNA-1 via microinjection into the lateral ventricle of mouse embryos, followed by IUE at E13. At E14, one day after IUE and following 6-hour pulse-labelling with EdU before sample collection, we observed a substantial reduction in the proportion of EdU+/GFP+ double-labelled cells in the shCdk5rap2 group 41.1 ± 3.1% as compared to the control group 58.9 ± 4.5% (Figure 7A-B). Remarkably, at E17, four days after IUE, Cdk5rap2 knockdown resulted in a significant number of GFP+ cells to persist in the VZ + SVZ (13.34 ± 3.31%) and IZ (27.85 ± 3.02%) with fewer GFP+ cells migrating towards the CP (58.81 ± 2.66%) as compared to control (Supplementary Figure 6H-I, Control, VZ + SVZ (4.20 ± 2.73%), IZ (14.29 ± 2.87%), CP (81.51 ± 3.26%). This data suggests that loss of Cdk5rap2 affects neuronal migration and proliferation *in vitro* and *in vivo*, similar to Arl2 loss-of-function.

**Fig. 7.**
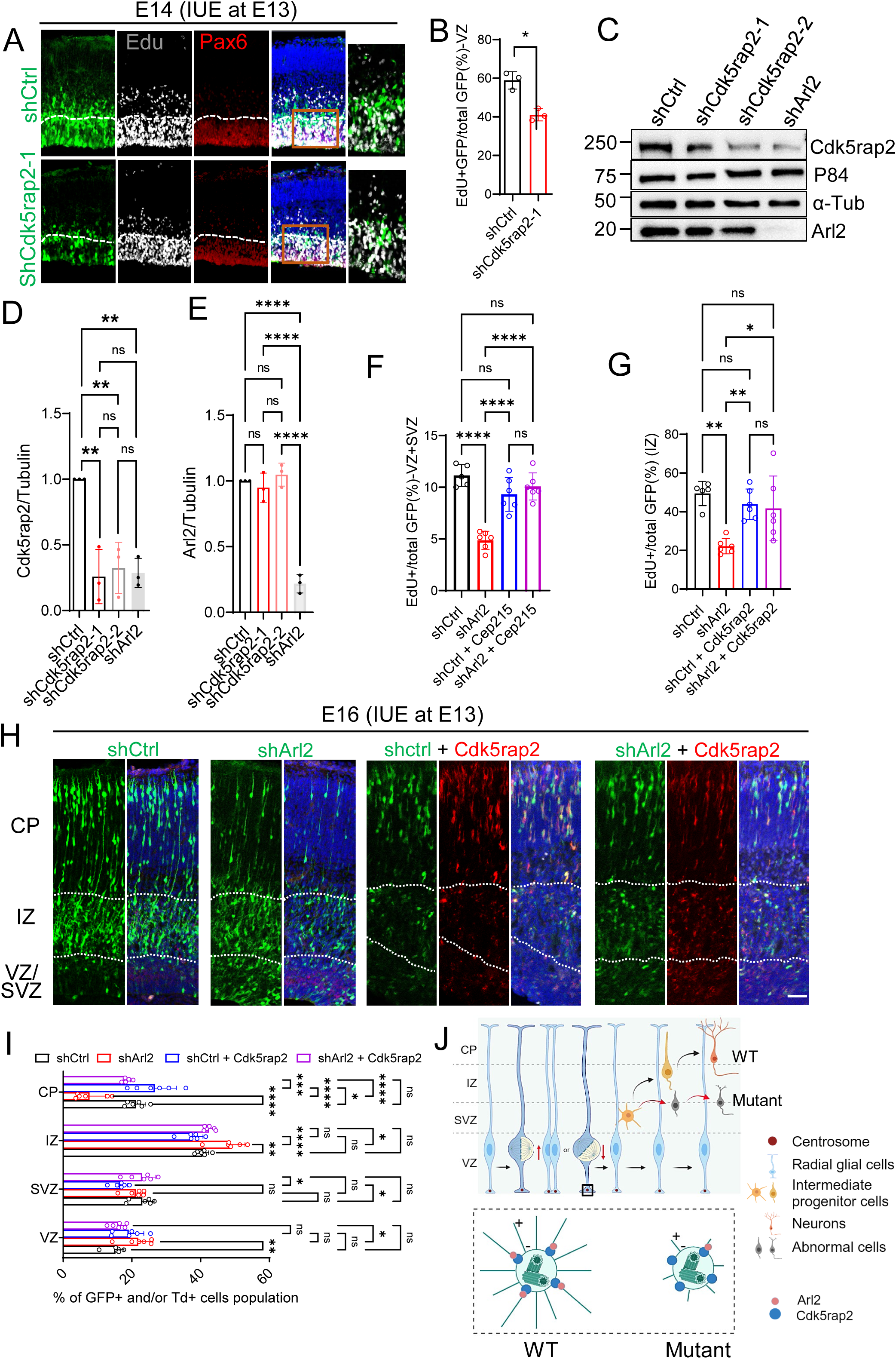
Arl2 functions upstream of Cdk5rap2 in regulating neuronal cell migration and proliferation in the developing cortex. (A) Brain slices from shCtrl (scrambled control) and shCdk5rap2 (Cdk5rap2 shRNA) groups at E14, one day after IUE, were labelled with EdU and GFP. (B) Bar graphs showing reduced EdU incorporation upon Cdk5rap2 knockdown (41.05 ± 3.16% in shCdk5rap2 vs 58.89 ± 4.50% in shCtrl). The values represent the mean ± s.d.. Student’s t-test in C, n = 3. Differences were considered significant at *p<0.05. (C) Western blotting analysis of mNPC protein extracts of control (H1-shctrl-GFP), Cdk5rap2 KD (H1-shCdk5rap2-GFP) and Arl2 KD with lentivirus (pPurGreen) infection in 48 h culture. Blots were probed with anti-Cdk5rap2 and anti-Arl2 antibody, α-Tubulin and p84 as loading control. (D) Bar graphs representing Cdk5rap2 protein levels upon Arl2 knockdown (shCdk5rap2-1 = 0.26 ± 0.21 and shCdk5rap2-2 = 0.32 ± 0.20; shArl2 = 0.29 ± 0.11 normalized in shCtrl, n = 3) in mNPCs. (E) Bar graphs representing Arl2 protein levels upon Cdk5rap2 knockdown (shCdk5rap2-1 = 0.95 ± 0.11 and shCdk5rap2-2 = 1.05 ± 0.09; shArl2 = 0.22 ± 0.07 normalized in shCtrl, n = 3) in mNPCs. (F) Bar graphs showing the total number of GFP+EdU+ double positive cells in VZ+SVZ by overexpression of Cdk5rap2 in Arl2 KD brains (Control = 11.14 ± 1.04%, shArl2 = 4.87 ± 0.81%, shCtrl + Cdk5rap2 = 9.32 ± 1.63%, shArl2 + Cdk5rap2 = 10.08 ± 1.31%) (shCtrl, n = 5 embryos; shArl2, shCtrl + shCdk5rap2, shArl2 + shCdk5rap2, n = 6 embryos) 2 days after IUE. (G) Bar graphs showing the total number of GFP+EdU+ double positive cells in IZ by overexpression of Cdk5rap2 in Arl2 KD brains (Control = 49.41 ± 6.25%, shArl2 = 22.17 ± 3.98%, shCtrl + Cdk5rap2 = 43.84 ± 7.89%, shArl2 + Cdk5rap2 = 41.68 ± 16.74%) (shCtrl, n = 5 embryos; shArl2, shCtrl + shCdk5rap2, shArl2 + shCdk5rap2, n = 6 embryos) 2 days after IUE. The values represent the mean ± s.d.. One-Way ANOVA in D-H. Differences were considered significant at ****p<0.0001. (H) Brain slices from shCtrl, shArl2, shCtrl + Cdk5rap2 and shArl2 + Cdk5rap2 groups at E16, three days after IUE, were labelled with GFP (shCtrl, shArl2, shCtrl + Cdk5rap2 and shArl2 + Cdk5rap2) and tdTomato (shCtrl + Cdk5rap2 and shArl2 + Cdk5rap2). (I) Bar graphs (images in H) representing GFP+ and/or Td+ cell population in the groups of control (VZ = 15.22 ± 2.47%, SVZ = 23.03 ± 3.46%, IZ = 40.55 ± 1.83%, CP=21.22 ± 2.82%, n = 6 embryos), shArl2 (VZ = 21.87 ± 4.31%, SVZ = 21.14 ± 2.87%, IZ = 48.45 ± 4.47%, CP = 7.70 ± 4.67%, n = 6 embryos), shCtrl + Cdk5rap2 (VZ = 19.08 ± 4.38%, SVZ = 16.59 ± 2.18%, IZ = 37.63 ± 3.05%, CP = 26.70 ± 6.21%, n = 6 embryos) and shArl2 + Cdk5rap2 (VZ = 16.07 ± 2.15%, SVZ = 22.98 ± 3.99%, IZ = 42.50 ± 1.45%, CP = 18.45 ± 1.51%, n = 6 embryos). (J) Working model (made by BioRender): Arl2 plays a novel role in regulating the balance of asymmetric and symmetric divisions of mNSCs and their proliferation and differentiation. Arl2 is required for the proliferation, migration and differentiation of mouse forebrain NPCs *in vitro* and *in vivo* by regulating centrosome assembly and microtubule growth in NPCs. Arl2 physically associates and recruits Cdk5rap2 to the centrosomes to promote microtubule assembly in NPCs. Arl2 functions upstream of Cdk5rap2 in regulating NPC proliferation and migration during mouse cortical development. The values represent the mean ± s.d.. Two-Way ANOVA with Multiple comparisons in I; Differences were considered significant at *p<0.05, **p<0.01, ***p<0.001 and ****p<0.0001, ns = non-significance. Scale bar; A and H = 50 µm.

### Arl2 is required for the centrosomal localization of Cdk5rap2 in mNPCs

Since we demonstrate that Arl2 co-localizes and physically associates with Cdk5rap2 at the centrosomes of mNPCs (Figure 6), we wondered whether Arl2 is required for the centrosomal localization of Cdk5rap2 in mNPCs. Indeed, Cdk5rap2 centrosomal localization was diminished upon Arl2 knockdown in mNPCs at interphase (supplementary Figure 6C-D) and mitosis (Figure 6B-C). Furthermore, Cdk5rap2 intensity were significantly reduced upon Arl2 knockdown in mNPCs at interphase (46.14 A.U.) (supplementary Figure 6C-D) and at metaphase (94.5 A.U.) (Figure 6B-C) as compared to the control (interphase: 77.37 A.U.; metaphase: 153.0 A.U.). Moreover, Cdk5rap2 protein levels were significantly reduced upon Arl2 knockdown in mNPCs by Western blotting analysis (Figure 7C-D; shCdk5rap2-1 = 0.26 ± 0.21 and shCdk5rap2-2 = 0.32 ± 0.20; shArl2 = 0.29 ± 0.11 normalized in shCtrl, n = 3). Conversely, Arl2 protein levels in mNPCs were not obviously affected by Cdk5rap2 knockdown as compared to the control (Figure 7C, E, shCdk5rap2-1 = 0.95 ± 0.11 and shCdk5rap2-2 = 1.05 ± 0.09; shArl2 = 0.22 ± 0.07 normalized in shCtrl, n = 3). Thus, Arl2 is required for localization and stabilization of Cdk5rap2 at the centrosomes in mNPCs.

### Cdk5rap2 overexpression rescues neurogenesis defects caused by Arl2 depletion in mouse developing cortex

To determine whether Cdk5rap2 is a physiological relevant target of Arl2 during neurogenesis *in vivo*, we overexpressed Cdk5rap2 along with Arl2 knockdown via microinjection into the lateral ventricle of mouse embryos, followed by IUE at E13, and examined cortical neurogenesis. Remarkably, Cdk5rap2 overexpression notably rescued neurogenesis defects caused by Arl2 KD *in vivo* (Figure 7). Two days after IUE at E15, the total number of GFP+EdU+ double positive cells in VZ+SZ as well as IZ was significantly rescued by overexpression of Cdk5rap2 in Arl2 KD brains (Figure 7F-G, VZ+SVZ; Control = 11.14 ± 1.04%, shArl2 = 4.87 ± 0.81%, shCtrl + Cdk5rap2 = 9.32 ± 1.63%, shArl2 + Cdk5rap2 = 10.08 ± 1.31%; IZ; Control = 49.41 ± 6.25%, shArl2 = 22.17 ± 3.98%, shCtrl + Cdk5rap2 = 43.84 ± 7.89%, shArl2 + Cdk5rap2 = 41.68 ± 16.74%). Furthermore, three days after IUE at E16, the number of GFP+ cells migrating to the CP in Arl2*-*depleted mouse brains by overexpressing Cdk5rap2 was dramatically increased to 18.45% compared with 7.70% in Arl2 KD brains (Figure 7H, I). These genetic data further support our model that Arl2 regulates NPC proliferation, migration and differentiation in mouse cortical development by interacting with Cdk5rap2 to promote microtubule growth from the centrosomes.

## Discussion

In this study, we demonstrate for the first time that the mammalian ADP ribosylation factor-like 2 (Arl2), a small GTPase, plays an important role in corticogenesis of the mouse brain. We have identified Arl2 as a new regulator in proliferation and differentiation of mouse NPCs and neuronal migration. Arl2 controls neurogenesis through the regulation of microtubule growth, independent of its function in mitochondrial fusion. We further demonstrate that Arl2 physically associates with Cdk5rap2, a centrosomal protein known to be important for microtubule organization and cortical development. Finally, Arl2 functions upstream of Cdk5rap2 to localize and stabilize Cdk5rap2 at the centrosome to regulate microtubule growth and neuronal migration (Figure 7J). Taken together, our data has identified a novel Arl2-Cdk5rap2 pathway in the regulation of microtubule growth and proliferation of mouse NPCs during cortical development.

### Arl2 regulates mouse corticogenesis via microtubule growth

Although mammalian Arl2 has been shown to be widely expressed in various tissues and is most abundant in the brain [30], the role of mammalian Arl2 in regulating mouse corticogenesis was unknown. In this study, we demonstrate the importance of Arl2 in regulating NPC proliferation and differentiation and neuronal migration during mouse cortical development. We provide evidence that Arl2 is required for centrosome assembly and spindle orientation in NPCs, similar to Cdk5rap2. Our finding is in line with the role of centrosomal/microtubule regulators in spindle orientation of NPCs [38, 39]. Our finding also suggest that mouse Arl2 has a novel role in neurite outgrowth in neurons *in vitro*. All these findings highlight the importance of mouse Arl2 in regulating microtubule growth during corticogenesis.

Similar to the previous finding that transgenic ARL2-Q70L animals exhibit reduced photoreceptor cell function and progressive rod degeneration [40], we found that overexpression of these mutant forms of Arl2 caused cell death of mouse NPCs both *in vitro* and *in vivo*. This is consistent with a recent report showing that lengthening mitosis of neural progenitor cells resulted in apoptosis of new-born neural progeny [41]. Likewise, human Arl2 plays an essential role for the survival of human embryonic stem cell-derived neural progenitor cells [42]. In human brain organoid models, defects in mitosis of neural stem cells is associated with decrease in stem cell number and apoptosis [18]. Given that our work highlights a novel role of mammalian Arl2 in mouse cortical development *in vivo* and the conservation of Arl2 in mouse and humans, it will be of great interest to investigate the role of human Arl2 in NPC divisions during cortical development.

It was reported that mitochondria functions are important for RG proliferation [43]. RGs display fused mitochondria, while new-born neurons have highly fragmented mitochondria right after mitosis of NPCs [43]. Increased mitochondria fission promotes neuronal fate, while induction of mitochondria fusion after mitosis redirect daughter cells toward self-renewal [43]. Although Arl2 is localized to mitochondria and regulates mitochondria fusion *in vitro* [26, 30, 32], we found that neither Arl2 knockdown nor overexpression of Arl2 obviously altered mitochondrial morphology in the mouse developing brain. In addition, overexpression of Arl^K71R^ mutant, which causes mitochondrial fragmentation without disrupting microtubule assembly [32], behaves similarly to Arl2 wild-type in the mNPC proliferation or neuronal migration during neurogenesis *in vivo* or in our microtubule regrowth assay *in vitro*. Therefore, the novel role of Arl2 in regulating neurogenesis in the developing cortex is most likely due to its role in microtubule growth, independent of its function in mitochondrial fusion.

### Arl2 plays a novel role in regulating neurogenesis via Cdk5rap2 function

Based on *in silico* analysis by AlphaFold multimer, co-immunoprecipitation, and PLA, we provide strong evidence that Arl2 physically associates with the centrosomal protein Cdk5rap2. Moreover, our super-resolution imaging clearly shows that Arl2 co-localizes with Cdk5rap2 at the PCM of the centrosomes. Cdk5rap2 is known to regulate centrosomal function and maintain the neural progenitor pool in the developing cortex [16, 44]. However, in addition to a similar defect in NPC proliferation upon Cdk5rap2 knockdown, we observed additional neuronal migration defects following Cdk5rap2 depletion that mimics Arl2 knockdown. Importantly, Cdk5rap2 overexpression rescues the loss of function phenotype of Arl2 in mice, leading to restored NPC proliferation and neuronal migration to the cortical plate. Therefore, Arl2 functions upstream of Cdk5rap2 in controlling NPC proliferation and neuronal migration via centrosomal functions. Pericentrin, another centrosomal protein required for cortical development, also interacts with and recruits Cdk5rap2 to the centrosome in the mNPCs [16]. Consistent with this finding, our analysis by AlphaFolder multimer also predicts that Arl2 can potentially interact with Pericentrin with an ipTM score of 0.33 (Figure 6E). The centrosome and the primary cilium at the apical RGs are intricately connected, both of which control NPC proliferation [45]. Interestingly, a recent study showed a role of Arl2 in cilia stability in rod photoreceptor neurons, as Arl2Q70L overexpression caused decreased function and degeneration of these cells [40]. Future study is warranted to determine whether Arl2 is also involved in ciliogenesis in RGs.

Radial glial cells exhibit a bipolar morphology with an apical process anchored to the ventricular surface and a basal process projecting towards the pial surface of the brain [46]. The centrosomes are located at the apical endfoot of the apical process, while microtubules in the basal process are largely acentrosomal where γ-tubulin was undetectable and instead are organized by Golgi outposts [47]. Whether Arl2 can also potentially involved in microtubule assembly within the basal process remains unknown and will be intriguing for future investigations.

Loss-of-function variants of Cdk5rap2 are associated with Primary autosomal-recessive microcephaly (MCPH) [48]. Although Arl2 variants have not been found in brain disorders, recent studies identified Arl2 as a candidate gene for an eye disorder [49] and its role in early photoreceptor development via its microtubule functions [50]. Interestingly, mouse Cdk5rap2 was also shown recently to be required for eye development by affecting retina progenitor cell proliferation and apoptosis [51], suggesting that Cdk5rap2 might be linked to Arl2 in other cell types beyond the developing cortex.

Taken together, our study highlights the critical role of Arl2 regulates NPC proliferation and neuronal migration during mouse cortical development. Mechanistically, Arl2 physically associates and recruits Cdk5rap2 to the centrosomes to promote microtubule assembly in NPCs and neuronal migration. These discoveries may facilitate the development of potential therapeutic strategies for neurodevelopmental disorders.

## Materials and Methods

### Animals

All animal studies were performed under the Institutional Animal care and use committee (IACUC) approved protocol (IACUC Protocol: 2016/SHS/1207 and 2021/SHS/1672).

C57BL/6 mice were purchased from InVivos for the in-utero electroporation and for primary mouse neural progenitor cell culture experiments.

### DNA constructs

Arl2 full length cDNA and 3 mutant form (Arl2^Q70L^, Arl2^T30N^ and Arl2^K71R^), and Cdk5rap2 from mouse and human were cloned into FUGW (Addgene plasmid # 14883) [52], FUtdTW (Addgene plasmid # 22478) [53], pBiFC-VC155 (Addgene plasmid # 22011) [54] and pBiFC-VN155 (I152L) (Addgene plasmid # 27097) [55] constructs. Small hairpin RNAs were cloned into pGreenPuro™ constructs from SBI, system biosciences (cat no: #s SI505A-1). Two shRNAs target different regions of mouse Arl2 (shArl2-1 and shArl2-2), Cdk5rap2 (shCdk5rap2-1 and shCdk5rap2-2) and one control shRNA with scrambled sequence were designed.

The following different sets of short hairpin sequences were cloned into pGreenPuro vectors: shArl2-1 (CATCGACTGGCTCCTTGATGACATTTCCA) and shArl2-2 (GACACTGGGCTTCAACATCAAGACCCTGG); shCdk5rap2-1 (GCACATCTACAAGACGAACAT) (Sigma, TRCN0000179786) and shCdk5rap2-2 (GCCATCAAGATACGATTCATT) (Sigma, TRCN0000183538).

### HEK293T culture and lentiviruses package

Clontech’s HEK 293T cell line were cultured in D-MEM high glucose medium (Invitrogen), containing 4.5 g/L D-glucose, and 4 mM L-glutamine. For packaging viral vector, high titers of engineered lentiviruses were produced by co-transfection of lentiviral vectors (FUGW, or FUtdTW or pGreenPuro), psPAX2 and pMD2.G into HEK293T cells followed by ultracentrifugation of viral supernatant as previously described [56].

### Mouse neural progenitor cells (mNPCs) culture

Mouse embryos were harvested at E14, and the dorsolateral cortex was dissected and enzymatically triturated to isolate NPCs as Supplementary Figure 1A. NPCs were suspension-cultured in Costar® 6-well Clear Flat Bottom Ultra-Low Attachment Multiple Well Plates (Corning) in proliferation medium (NeuroCult™ Proliferation Kit (Mouse & Rat), STEMCELL) containing human EGF (10 ng/ml), human FGF2 (10 ng/ml) (Invitrogen, Carlsbad, CA), N2 supplement (1%) (GIBCO), penicillin (100 U/ml), streptomycin (100 mg/ml), and L-glutamine (2 mM) for 7 days and were allowed to proliferate to form neurospheres. DIV 7 neurospheres were dissociated into single cells using accutase, yielding 5-6 x 10^6^ cells per 6-well plate. For proliferation assay, forty-eight hours lentivirus (pPurGreen) infection, the cells were pulsed with 1 mM 5-Ethynyl-2’-deoxyuridine (EdU, Invitrogen) for 3 hr. In vitro NPC differentiation assay, mNPC cells were seeded onto 24-well plate with 60 mm coverslips coated with poly-L-lysine, at a density of 4.5 x 10^4^ cells/coverslip. Twenty-four hours lentivirus (pPurGreen) infection, NPCs were cultured as monolayer in differentiation medium containing B27 (2%) in Neurobasal medium and were maintained for 5–6 days.

### Cortical primary neuron culture

Primary cultures of cortical neurons were prepared from embryonic day 18 (E18) mice as previously described [56]. Briefly, the cortex was carefully dissected from the E18 brain in Earle’s Balanced salt solution (EBSS – Gibco 0766) and collected in buffer (127 mM NaCl, 5 mM KCl, 170 μM Na2HPO4, 205 μM KH2PO4, 5 mM Glucose, 59 mM Sucrose, 100 U/mL Penicillin/Streptomycin, pH 7.4). Cells were dissociated using 25 mg/ml papain. After collection in growth medium (Dulbecco’s Modified Eagle’s Medium w/GlutaMax (Invitrogen) containing 1 M HEPES, 10% heat inactivated Horse Serum (Invitrogen), and 100 U/mL Penicillin/Streptomycin, pH 7.4) cells were filtered through a 70 µM cell strainer. Subsequently, cells were seeded onto 24-well plate with 60 mm coverslips coated with poly-L-lysine, at a density of 4.5 x 10^4^ cells/coverslip.

### In-utero electroporation

In-utero electroporation was performed as described previously [57]. Pregnant E13 mice were anesthetized with isoflurane and proceeded with the laparotomy procedure. Small hairpin plasmid DNA with the GFP or overexpression plasmid with tdTomato reporter (2-3 µg/µl) was injected into the lateral ventricles of the embryos through the uterine wall. Subsequently for the electroporation, four electrical pulses of 35V, 50 msec was administered with the electroporator device and the mice were allowed to undergo normal development after the surgery. The electroporated embryonic mice brains were harvested at E14, E15, E16 and E17 for the cell proliferation, differentiation and migration analysis.

### EdU (5-Ethynyl-2’-deoxyuridine) incorporation assay

For EdU labelling experiments in mice, EdU was injected intraperitonially into the pregnant mice and the mice were sacrificed after 6hrs for brain harvest. The brain samples were subjected to standard immunochemistry procedure. The incorporated EdU was detected using fluro azide from ClickiT® EdU Imaging Kit (Invitrogen).

### Tissue preparation and immunostaining analysis

Embryonic mice were dissected in PBS and the embryonic brain samples were fixed in 4% Paraformaldehyde overnight, subsequently the brain samples were stored in 30% sucrose prior to sectioning. The brain samples were mounted in Tissue-Tek embedding medium and were sectioned using cryostat. For mouse neural progenitor cells, the cells were grown on coverslips and fixed with 4% Paraformaldehyde for 15 mins at RT. Subsequently the cells were washed with PBS twice and stored prior to staining. Cells and brain sections were consequently washed with TBS and blocked with 5% Normal donkey serum in TBS with 0.1% Triton X (TBST). Respective primary antibodies were prepared with the blocking solution and incubated for 2 hours at RT. Subsequently the cells were washed with TBST and proceeded with secondary fluorophore antibodies incubation for 1 hour at RT. For the detection of EdU incorporated cells and tissues, Alexa fluor azide was used as per the protocol described (ClickiT® EdU Imaging Kits; Invitrogen). The cells and tissues were washed and mounted for imaging. Micrographs were taken using LSM710 confocal microscope system (Axio Observer Z1; ZEISS), fitted with a PlanApochromat 40x/1.3 NA oil differential interference contrast objective, and brightness and contrast were adjusted by ImageJ.

The primary antibodies used in this paper, rabbit anti-TBR1 (1:500; Cell signalling, cat no: 49661S), rabbit anti-TBR2 (1:500; abcam, cat no: ab23345), rabbit anti-Pax6 (1:500; BioLegend, cat no: B328397), mouse anti-alpha tubulin (1:1000; Sigma, cat no: T6199), mouse anti-gamma tubulin (1:500; Sigma, cat no: T5326), mouse anti-DCX (1:300; Cruz biotechnology, cat no: A0919), rabbit anti-Ki67 (1:500; abcam, cat no: ab16667), rat anti-PH3 (1:500; sigma, cat no: 4882), rabbit anti-NeuroD2 (1:500, abcam, cat no: ab104430), rabbit anti-caspase3 (1:500; BD Pharmingen, cat no: 559569), rabbit anti-Cdk5rap2 (1:500, Merck Millipore, cat no: 06-1398), rabbit anti-Arl2 (1:300; abcam, cat no: ab183510), mouse anti-myc (1:500; abcam, cat no:1011022-5), guinea pig anti-GFP (1:1000; Dr. Yu Feng Wei lab), mouse anti-GFP (1:1000; Dr. Yu Feng Wei lab), rabbit anti-HA (1:500; Sigma, cat no: H6908), rat anti-HA (1:500; Roche, cat no:423001).

### Proximity ligation assay

Proximity ligation assay (PLA) was performed as described ([58], Adopted from Duolink PLA, Merck). Mouse neural progenitor cells were transfected with the following constructs including control-GFP, control-myc, Arl2-GFP and Cdk5rap2-myc using Lipofectamine Transfection reagent (Invitrogen). The cells were washed with cold PBS thrice and fixed with 4% paraformaldehyde in PB for 15 min. Subsequently, the cells were blocked with 5% normal donkey serum in TBS-Tx (0.1% Triton-X100) for 45 min. The cells were incubated with respective primary antibodies at RT for 2 hrs. The cells were then incubated with PLA probes at 37°C for 1 hr. Subsequently, the cells were washed Buffer A for 5mins at RT. The cells were proceeded with ligation of probes at 37°C for 30 min and amplification at 37°C for 1.5 hrs, followed by two washes with Buffer B at RT. The cells were washed once with 0.01x Buffer B and proceeded with primary antibodies incubation diluted in 3% BSA in PBS for 2 hrs at RT. Following this, the cells were washed twice with 0.1% TBS-TX and incubated with secondary antibodies for 1.5 hrs at RT. The cells were subsequently washed with PB and then mounted using *in situ* mounting media with DAPI (Duolink, Sigma-Aldrich).

### Microtubule regrowth assay

Mouse neural progenitor cells were incubated with 2.5mM thymidine at 37°C for 20 hrs to induce S phase arrest (Figure 3A). Subsequently the cells were released from S phase arrest for 7 hrs. The cells were then incubated with 200 nM nocodazole for 4 hrs. The cells were washed with ice cold medium and incubated on ice for 30 min to initiate microtubule depolymerisation. The cells were subsequently replaced with pre-warmed medium at 37°C. The cells were washed with PBS and were incubated in 4% paraformaldehyde for 15 min to fix the cells. The standard immunochemistry assay was performed to quantify the Alpha-tubulin density.

### Co-immunoprecipitation

Cells were lysed using PierceTM IP lysis buffer (ThermoFisher Scientific, cat no: 87787) with protease inhibitors. 1% cell lysate was taken for input controls and the remaining were incubated with respective pulldown antibodies overnight at 4°C. Protein A/G ultra-link resin beads (ThermoFisher Scientific, cat no: 53135) were added to the cell lysate and incubated for 3 hours at 4°C. Consequently, the beads were washed with PBS for several times to remove the residual proteins. The beads were mixed with the protein loading dye and proceeded for western blot analysis.

### Western blot analysis

Cells were lysed using PierceTM RIPA buffer (ThermoFisher Scientific, cat no: 89901) with protease inhibitors. The proteins samples were separated using SDS-PAGE and were transferred onto the nitrocellulose membrane. The membranes were blocked with low fat dry milk in Phosphate buffered saline (PBS) with 0.1% tween20 (PBST) for 1 hr at RT. Subsequently, the membranes were incubated with respective primary antibodies in 5% Bovine serum albumin (BSA) with PBST overnight at 4°C. The membranes were washed thrice with PBST and incubated with HRP-conjugated secondary antibodies to probe the target proteins for 1 hr at RT. The membranes were washed and the proteins were detected using Super signalTM west pico chemiluminescence substrate (Protein biology, cat no: 34580).

### AlphaFold2-multimer protein complex prediction

To discover Arl2 interactors with centrosome proteins, we performed protein complex predictions using Alphafold-multimer developed by DeepMind [59]. Arl2 was predicted against core centrosome proteins. All the predictions were performed using AlphaPulldown Pipeline v0.30.6 [35] with default settings. Multiple sequence alignments (MSAs) and template input to the Alphafold-multimer were calculated by MMseqs2 [36]. To analyse the results produced by AlphaFold-multimer, interface pTM (ipTM) scores from the predictions were used to evaluate the interaction possibility and confidence. Predicted interaction structure model were used for further analysis.

### Spinning disc super-resolution imaging

Super-resolution imaging was performed as previously described [60]. In brief, super-resolution Spinning Disc Confocal-Structured Illumination Microscopy (SDC-SIM) was performed on a spinning disk system (Gataca Systems) based on an inverted microscope (Nikon Ti2-E; Nikon) equipped with a confocal spinning head (CSU-W; Yokogawa), a Plan-Apo objective (100×1.45-NA), a back-illuminated sCMOS camera (Prime95B; Teledyne Photometrics) and a super-resolution module (Live-SR; GATACA Systems). All image acquisition and processing were controlled by the MetaMorph (Molecular Device) software. Images were further processed with imageJ.

### Live-cell imaging

To capture time-lapse images of mouse neural progenitor cells (mNPCs), a super-resolution spinning disk confocal-structured illumination microscopy equipped with a Plan-Apo objective (100× 1.45-NA) was used. The imaging was conducted in a chamber at a temperature of 37°C with CO2 supplement. mNPCs were imaged for 16 hours (5 min each time interval for Figure 4H and Sup. Figure 5A) or for 5 min without intervals (for Figure 5C). The videos were processed using and ImageJ software. Mito-RFP tracker (Plasmid #51013) was from Addgene (pLenti.CAG.H2B-cerFP-2A-mito-dsRFP.W). Viafluor-488 live cell microtubule staining kit (Biotium, #70062) were used for live imaging of mNPCs *in vitro*.

### Tracking of EB3-GFP or EB3-Td comets

mNPCs expressing EB3-GFP or EB3-Td were subjected to live-cell imaging using a super-resolution spinning disk confocal-structured illumination microscopy as mentioned above. The amount and velocity of the EB3-GFP comets were calculated and kymographs were generated using KymoButler [61]. A cell was imaged for 3-5 min without time interval for each movie and videos were generated with NIH ImageJ software.

### Statistical analysis

All experiments were repeated at least thrice, and comparable results were obtained. All statistical analysis was performed using GraphPad prism. Paired or unpaired T-test were used for the comparison of two independent groups and one-way or two-way Anova were used for comparing more than two independent groups. Statistical significance was represented by ***p <0.001, **p<0.01, *p<0.05 compared with the control groups.

## Acknowledgments

This work is supported by Singapore Ministry of Education Tier 2 MOE-T2EP30121-0002 to H.W.

## Author contributions

Conceptualization, DM and HW; Methodology, Data curation, and formal analysis, DM, K-YL, SD, JL, MG, YST, HYA, ASV, GY, TC; Writing-original draft, DM, SD, MG and HW; Writing-review & editing, HW, DM, and MG; funding acquisition, HW; Resources, HW; Supervision, DM and HW.

## DECLARATION OF INTERESTS

The authors declare no competing interests.

## Figure legends

**Supp Figure 1.**
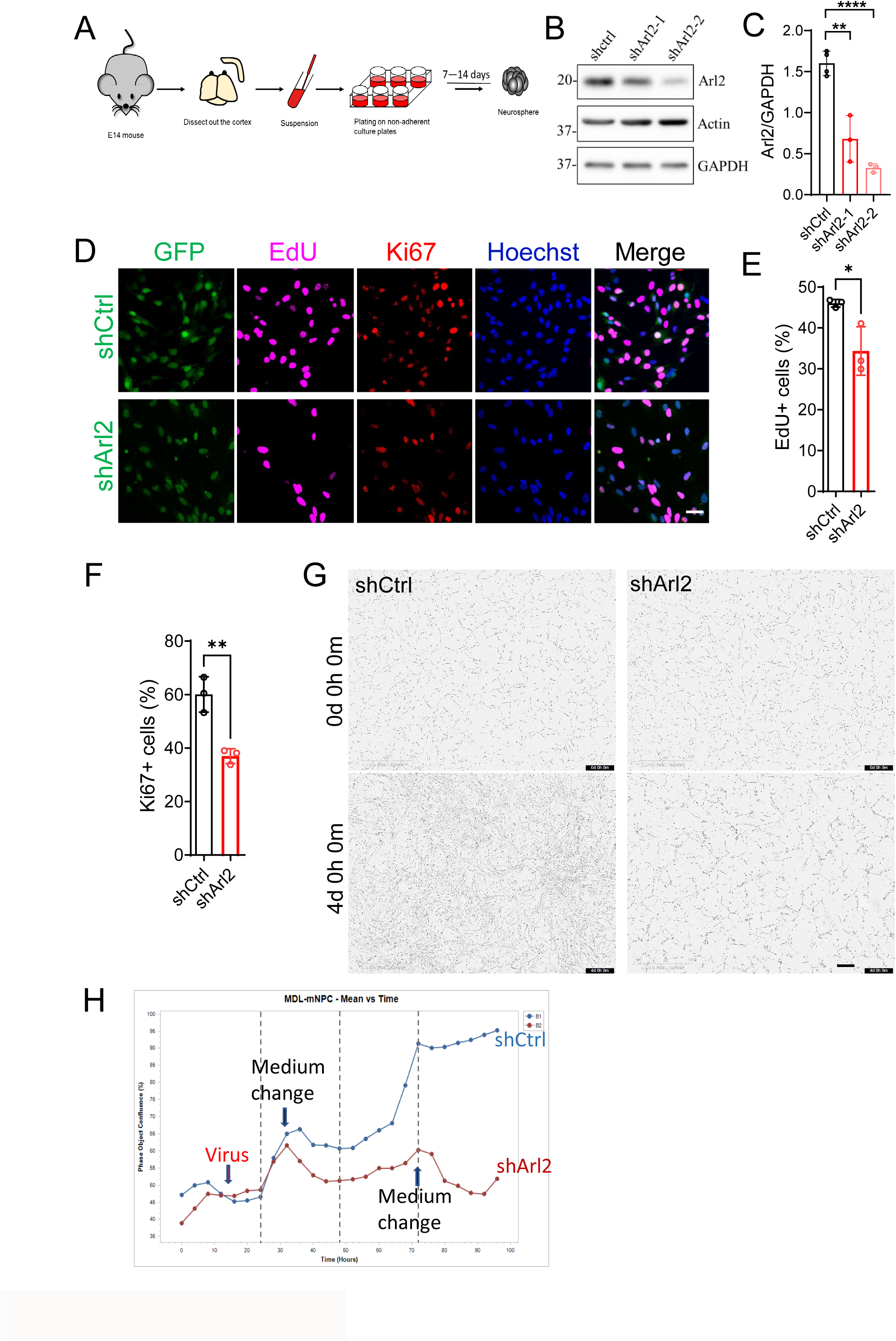
Arl2 knockdown affects mouse neural progenitor cell proliferation in vitro. (A) Schematic representation of primary mouse neural progenitor cell (mNPC) culture from embryonic mice. (B) Western blotting analysis of mNPC protein extracts of control (H1-shctrl-GFP) and Arl2 knockdown (H1-shArl2-GFP) with lentivirus (pPurGreen) infection in 48 h culture. Blots were probed with anti-Arl2 antibody and anti-GAPDH antibody. Immunoblot showing the efficiency of Arl2 knockdown. (C) Bar graphs representing knockdown efficiency of Arl2 normalized to the internal control GAPDH (0.68 ± 0.28 in shArl2-1, 0.33 ± 0.05 in shArl2-2 vs 1.60 ± 0.15 in shctrl, n = 3). (D) Immunostaining micrographs of mNPCs labelled for EdU and the proliferation marker Ki67. (E) Bar graphs showing reduced EdU incorporation upon Arl2 knockdown (34.35 ± 5.95% in shArl2 with 12 images vs 46.03 ± 0.95% in shctrl with 14 images, n = 3 batches). (F) Bar graphs showing decreased Ki67+ cells in shArl2 group compared to the control (36.98 ± 2.77% in shArl2 with 12 images vs 60.09 ± 6.63% in shCtrl with 14 images, n = 3 batches). (G) Time series showing decreased cell proliferation upon Arl2 knockdown as compared to control. (H) Line graph representing the timeline of mNPC proliferation and showing the defects in cell proliferation in shArl2 group compared to the control. At least three sets of independent experiments were performed. The values represent the mean ± s.d.. One-Way ANOVA in C. Student’s t-test in E, F. Differences were considered significant at at *p<0.05, **p<0.01 and ****p<0.0001. Scale bars; D = 30 µm; G= 250 µm.

**Supp Figure 2.**
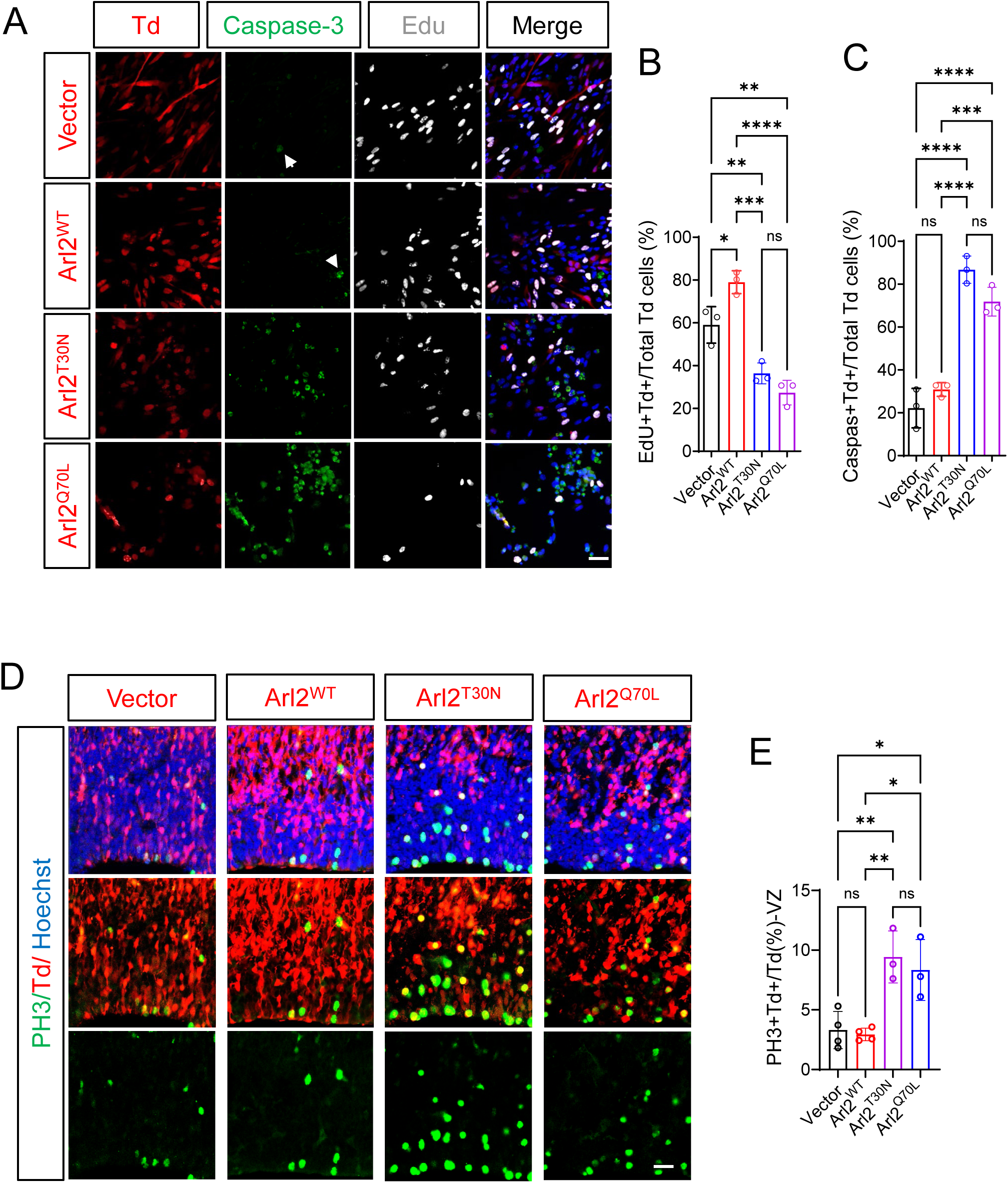
Overexpression Arl2 and mutant affect mouse neural progenitor cell proliferation. (A) Immunostaining micrographs of mNPCs *in vitro* in Arl2^WT^, Arl2^T30N^ and Arl2^Q70L^ labelled with EdU and caspase-3. (B) Bar graphs showing the proportion of EdU+ cells in control (59.06 ± 8.52%), Arl2^WT^ (79.01 ± 5.34%), Arl2^T30N^ and Arl2^Q70L^ (36.4 ± 4.93% and 27.4 ± 5.78%, respectively) (n = 3). (C) Bar graphs showing the proportion of caspase-3 + cells in control (22.18 ± 9.21%, n = 4), Arl2^WT^ (30.89 ± 3.22%, n = 4), Arl2^T30N^ and Arl2^Q70L^ (86.76 ± 6.36% and 71.86 ± 6.74%, respectively) (n = 3). (D) Brain slices from Arl2^WT^, Arl2^T30N^ and Arl2^Q70L^, at E15, two days after IUE, labelled for phospho-histone H3-positive (PH3+). (E) Bar graphs showing the proportion of PH3+ cells in control 3.31 ± 1.56%, n = 4; Arl2^WT^ 2.94 ± 0.53%, n = 4; Arl2^T30N^ 8.33 ± 2.55% and Arl2^Q70L^ 9.43 ± 2.18%, n = 3, in two day after IUE in VZ of brain sections. The values represent the mean ± s.d.. One-Way ANOVA with Multiple comparisons in B, C and E. Differences were considered significant at at *p<0.05, **p<0.01, ***p<0.001 and ****p<0.0001, ns = non-significance. Scale bars; A and D = 50 µm.

**Supp Figure 3.**
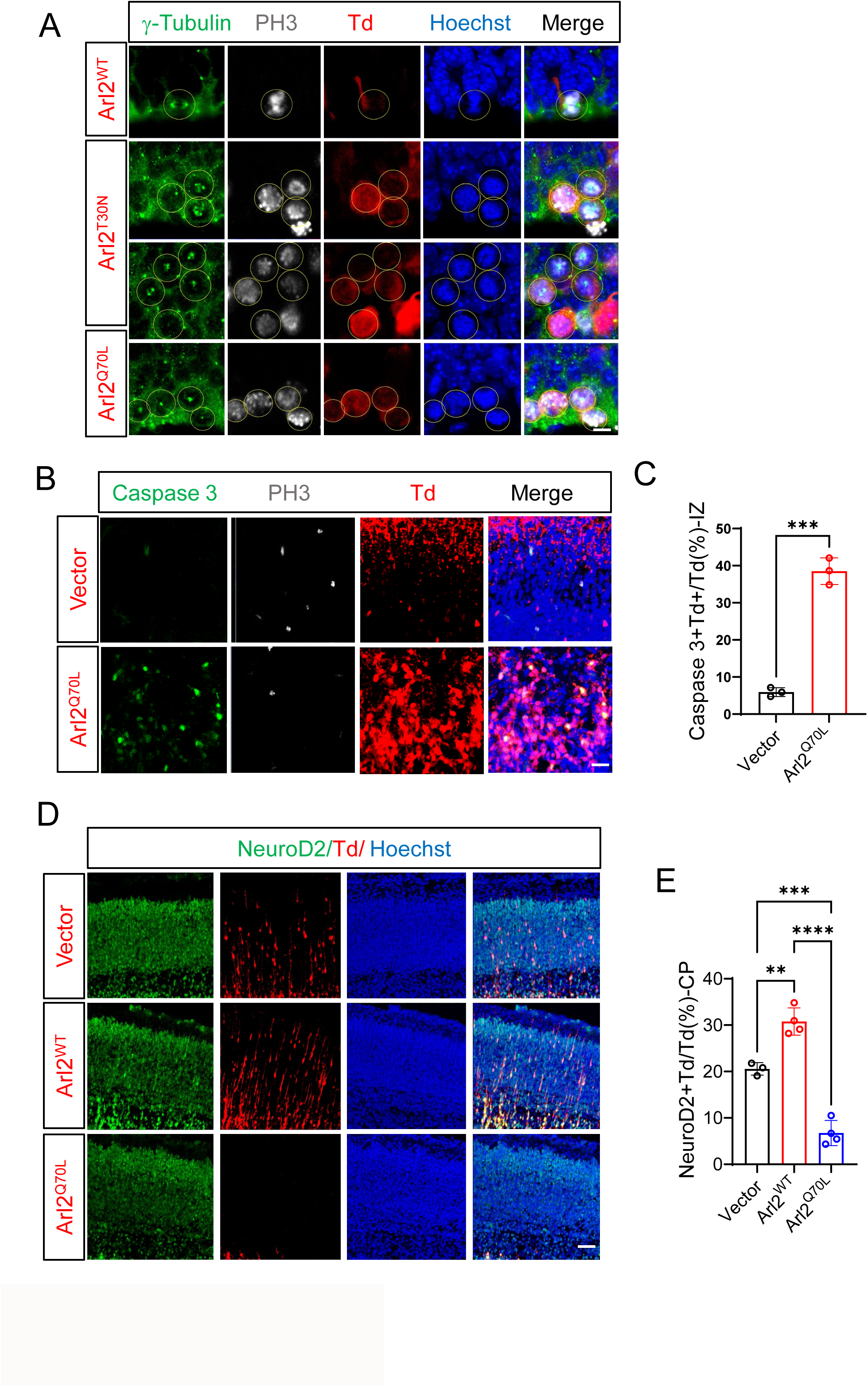
Overexpression Arl2 mutant affect mouse neural progenitor cell proliferation. (A) Immunostaining micrographs of mNPCs *in vitro* in Arl2^WT^, Arl2^T30N^ and Arl2^Q70L^ labelled for phospho-histone H3-positive (PH3+). (B) Brain slices from Arl2^WT^ and Arl2^Q70L^, at E16, three days after IUE, labelled for caspase-3, tdTomato and PH3. (C) Bar graph showing the caspase-3 staining and the proportion of caspase-3 + cells in the IZ in Arl2^Q70L^ (38.52 ± 3.60%) as compared to control (5.93 ± 1.19%) (n = 3) in 3 days after IUE. (D) Brain slices from Vector control, Arl2^WT^ and Arl2^Q70L^, at E16, three days after IUE, labelled for Neuro-D, tdTomato and Hoechst. (E) Bar graph showing the expression of NeuroD2, a neuronal marker found in immature neurons, in Arl2^WT^ (30.77 ± 2.93%) but dramatically reduced in Arl2^Q70L^ (6.75 ± 2.69%) 3 days after IUE as compared to control (20.57 ± 1.36%). The values represent the mean ± s.d.. One-Way ANOVA in E. Student’s t-test in C, Differences were considered significant at **p<0.01, ***p<0.001 and ****p<0.0001, ns = non-significance. Scale bars; A = 5 µm; B = 50 µm; C = 80 µm.

**Supp Figure 4.**
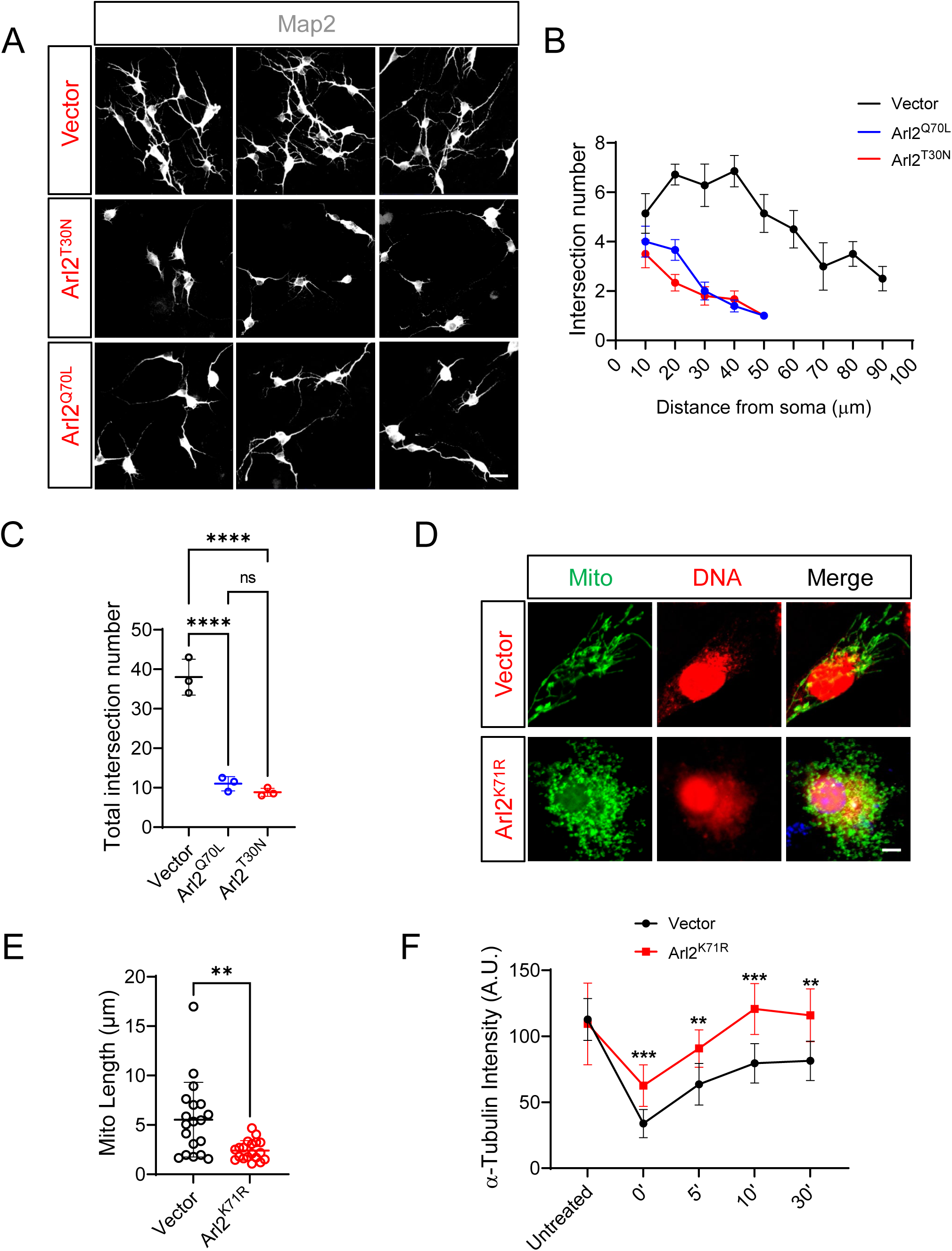
Arl2-K71R mutant causes mitochondrial fragmentation and has no effect on microtubule regrowth. (A) Immunostaining micrographs showing decreased neural complexities of in primary cortical neurons (labeled by Map2) *in vitro* in overexpression of Arl2^T30N^ and Arl2^Q70L^ mutants. (B) Scholl’s analysis showing the distance from soma and the intersection number reduced in the ARL2^T30N^ and ARL2^Q70L^ mutants as compared to the control. (C) Scholl’s analysis showing the total intersection number as measured and significantly reduced in Arl2^Q71L^ (11 ± 1.80) and Arl2^T30N^ (8.83 ± 1.04) as compared to control (38 ± 4.58). (D) Immunostaining micrographs of mNPCs in Vector control and mouse Arl2^K71R^ overexpression. Arl2^K71R^ overexpression showed fragmented mitochondria with shortened mitochondrial length as compared to control in mNPCs *in vitro*. (E) Graph representing mitochondrial length (2.41 ± 0.99 µm) in Arl2^K71R^ mutant as compared to control (5.53 ± 3.78 µm). (F) Line graph of microtubule regrowth assay representing α-tubulin intensity in overexpression of Arl2^K71R^ mutant group (Untreated =109.41 ± 30.92, 0 min = 62.56 ± 15.69, 5 min = 90.77 ± 14.18, 10 min = 120.69 ± 19.26, and 30 min =115.93 ± 20.06) as compared to the control (Untreated = 112.73 ± 15.82, 0 min = 33.82 ± 10.65, 5 min = 63.64 ± 15.80, 10 min = 79.52 ± 14.88, and 30 min = 81.37 ± 14.90) (Unit = A.U.) in mNPCs in vitro. The values represent the mean ± s.d.. One-Way ANOVA in C, Student’s t-test in E, Multiple t-test in F, n = 3. Differences were considered significant at *p<0.05, **p<0.01, ***p<0.001 and ****p<0.0001, ns = non-significance. Scale bars; A = 20 µm; D = 5 µm.

**Supp Figure 5.**
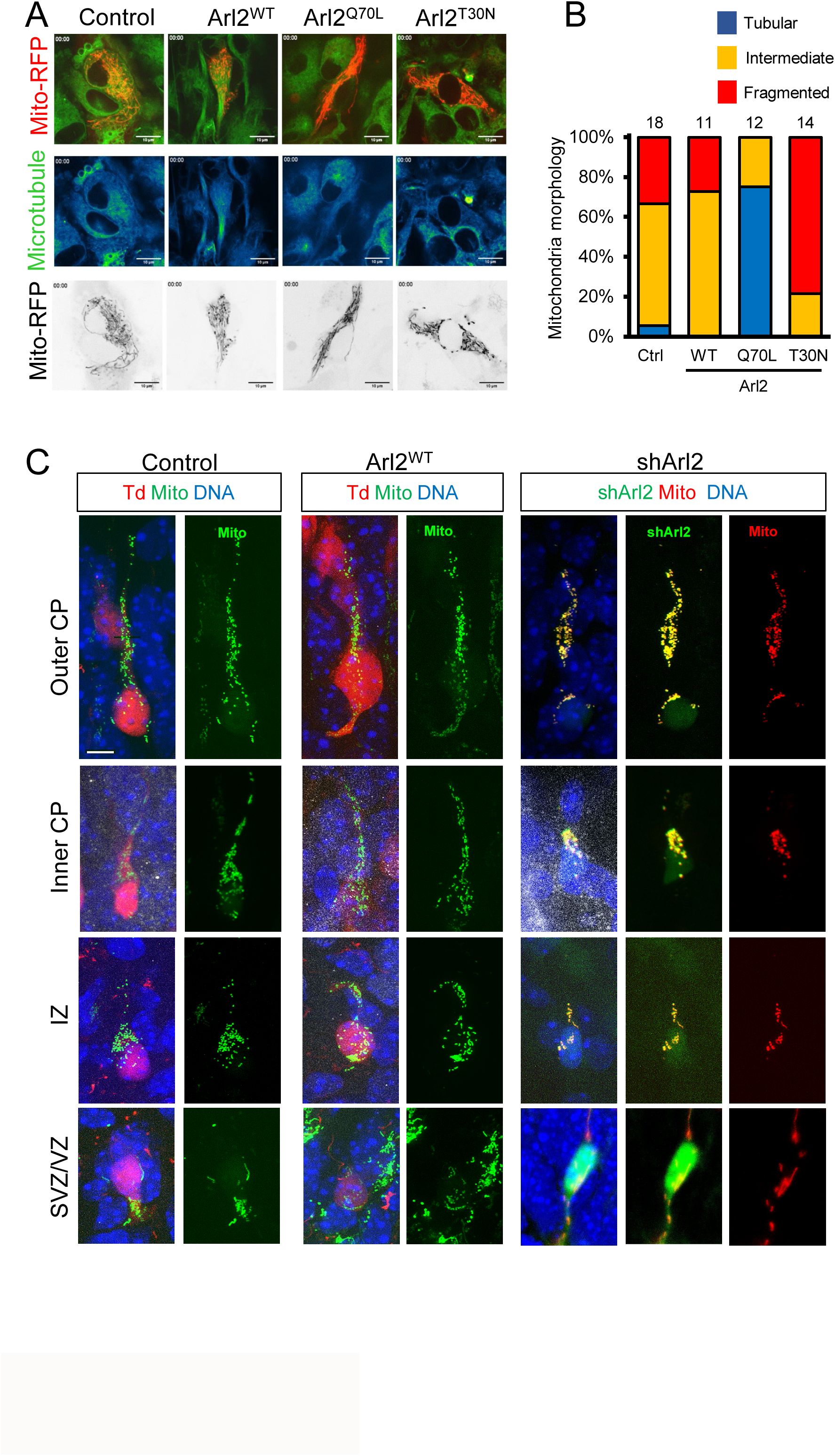
Mitochondrial morphology remained unchanged upon shArl2 KD or overexpression of Arl2 in mouse brains. (A) Live imaging micrograph of mitochondria (Mito-RFP, Plasmid #51013, Addgene) and microtubule (Viafluor-488 live cell microtubule staining kit (Biotium, #70062) dynamics in control, Arl2^Q70L^-overexpressing, or Arl2^T30N^-overexpressing mNPCs. (B) Bar graph shows qualifications of mitochondria morphology in various genotypes in A. (C) Brain slices from shCtrl, Arl2^WT^ and shArl2 groups at E17, four days after IUE, were labelled with GFP, TdTomato, Mito and DNA. Scale bars; A and C = 10 µm.

**Supp Figure 6.**
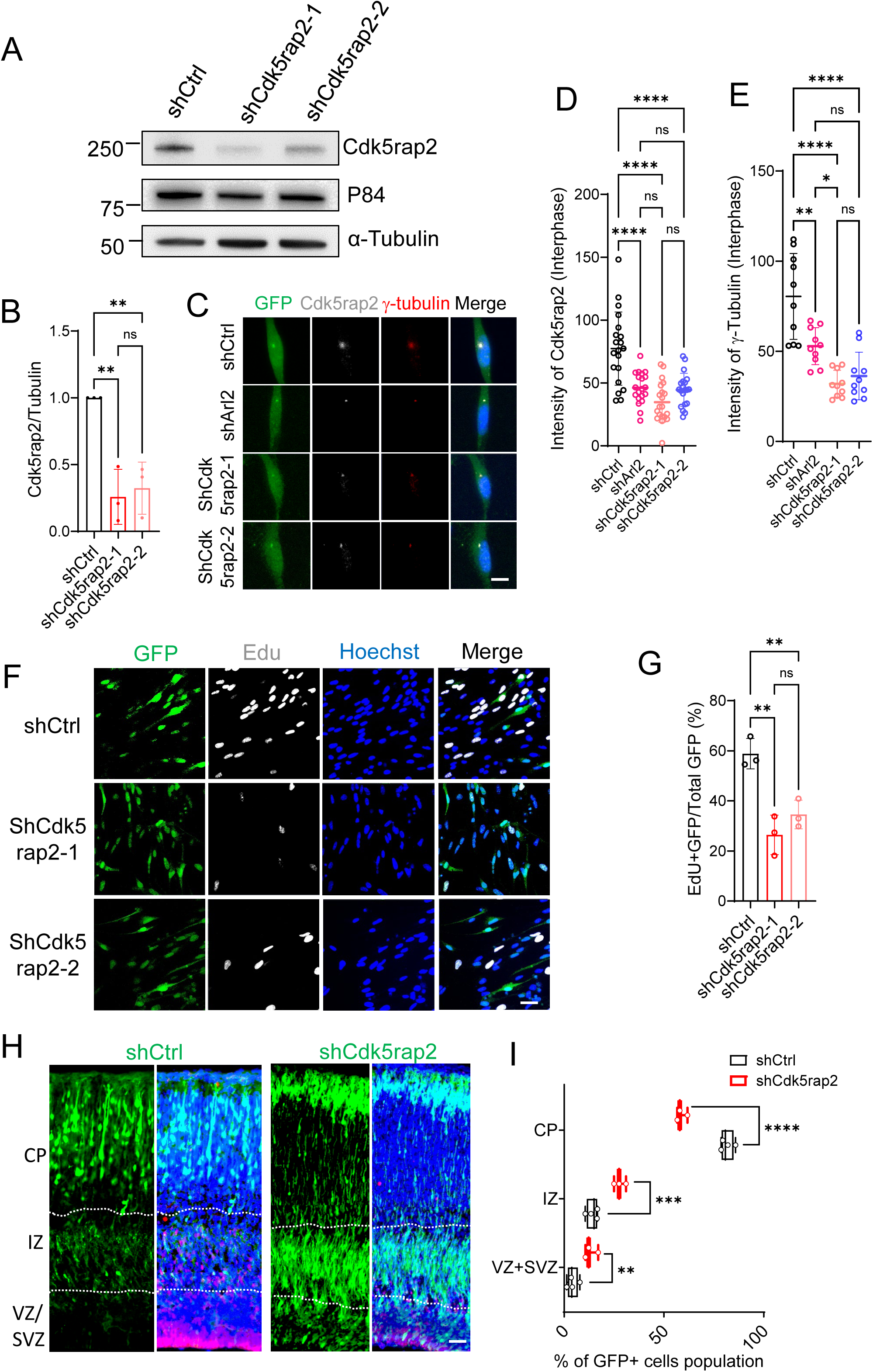
The phenotypes of Cdk5rap2 knockdown are similar with Arl2 KD. (A) Western blotting analysis of mNPC protein extracts of control (H1-shctrl-GFP) and Cdk5rap2 knockdown (H1-shCdk5rap2-GFP) with lentivirus (pPurGreen) infection in 48 h culture. Blots were probed with anti-Cdk5rap2 antibody and anti-GAPDH antibody. (B) Bar graphs representing knockdown efficiency of Cdk5rap2 normalized to the internal control GAPDH (0.26 ± 0.21 in shCdk5rap2-1, 0.32 ± 0.20 in shCdk5rap2-2 normalized in shCtrl, n = 3). (C) Immunostaining micrographs in shCtrl, shArl2, shCdk5rap2-1 and shCdk5rap2-2 in interphase mNPCs were labelled with γ-tubulin, Cdk5rap2 and Hoechst (DNA). (D) Quantification graph showing Cdk5rap2 intensity at interphase of mNPCs (46.14 ± 12.51) upon Arl2 knockdown and (shCdk5rap2-1 = 34.73 ± 15.93; shCdk5rap2-2 = 44.53 ± 13.16) upon Cdk5rap2 knockdown as compared to shCtrl (77.37 ± 29.09, n = 3 batches with 20 cells). (E) Quantification graph showing γ-tubulin intensity at interphase of mNPCs (52.83 ± 10.22) upon Arl2 knockdown and (shCdk5rap2-1 = 32.02 ± 7.45; shCdk5rap2-2 = 36.29 ± 13.21) upon Cdk5rap2 knockdown as compared to shCtrl (80.47 ± 23.81, n = 3 batches with 10 cells). The values represent the mean ± s.d.. One-Way ANOVA in D and E. Differences were considered significant at *p<0.05, **p<0.01 and ****p<0.0001, ns = non-significance. (F) Immunostaining micrographs in shCtrl, shCdk5rap2-1 and shCdk5rap2-2 in mNPCs were labelled with EdU and DNA. (G) Bar graphs showing reduced EdU incorporation upon Cdk5rap2 KD in mNPCs. The values represent the mean ± s.d. (shCtrl = 58.84 ± 6.06%; shCdk5rap2-1 = 26.42 ± 7.89%; shCdk5rap2-2 = 34.58 ± 5.72%, n = 3). (H) Brain slices from shCtrl, shCdk5rap2 groups at E17, four days after IUE, were labelled with GFP. (I) Box plots representing GFP+ cells for (H) in CP (shctrl: 81.51 ± 3.26%, shCdk5rap2: 58.81 ± 2.66%), IZ (shctrl: 14.29 ± 2.87%, shCdk5rap2: 27.85 ± 3.02%), and SVZ + VZ (shctrl: 4.20 ± 2.73%, shCdk5rap2: 13.34 ± 3.31%) (shCtrl: n = 4, shCdk5rap2: n = 3) showing defects in neuronal migration to CP upon Cdk5rap2 Knockdown compared to the control. The values represent the mean ± s.d.. One-Way ANOVA in B, D, E and G. Multiple unpaired t tests in I. Differences were considered significant at **p<0.01, ***p<0.001 and ****p<0.0001. ns = non-significance. Scale bars; C = 5 µm; F = 50 µm; H = 80 µm.

## Notes

### Competing Interest Statement

The authors have declared no competing interest.

## References

1. Taverna E, Gotz M, Huttner WB. The cell biology of neurogenesis: toward an understanding of the development and evolution of the neocortex. Annu Rev Cell Dev Biol. 2014;30:465–502. Epub 20140627. doi: 10.1146/annurev-cellbio-101011-155801. PubMed PMID: 25000993.

2. Shitamukai A, Matsuzaki F. Control of asymmetric cell division of mammalian neural progenitors. Development, growth & differentiation. 2012;54(3):277–86.

3. Itoh Y, Tyssowski K, Gotoh Y. Transcriptional coupling of neuronal fate commitment and the onset of migration. Current Opinion in Neurobiology. 2013;23(6):957–64. doi: 10.1016/j.conb.2013.08.003.

4. Furutachi S, Miya H, Watanabe T, Kawai H, Yamasaki N, Harada Y, et al. Slowly dividing neural progenitors are an embryonic origin of adult neural stem cells. Nature Neuroscience. 2015;18(5):657–65. doi: 10.1038/nn.3989.

5. Telley L, Agirman G, Prados J, Amberg N, Fièvre S, Oberst P, et al. Temporal patterning of apical progenitors and their daughter neurons in the developing neocortex. Science. 2019;364(6440):eaav2522. doi: doi:10.1126/science.aav2522.

6. Beattie R, Hippenmeyer S. Mechanisms of radial glia progenitor cell lineage progression. FEBS Lett. 2017;591(24):3993–4008. Epub 20171122. doi: 10.1002/1873-3468.12906. PubMed PMID: 29121403; PubMed Central PMCID: PMCPMC5765500.

7. Dwyer ND, Chen B, Chou S-J, Hippenmeyer S, Nguyen L, Ghashghaei HT. Neural Stem Cells to Cerebral Cortex: Emerging Mechanisms Regulating Progenitor Behavior and Productivity. The Journal of Neuroscience. 2016;36(45):11394–401. doi: 10.1523/jneurosci.2359-16.2016.

8. Shahbazi MN, Zernicka-Goetz M. Deconstructing and reconstructing the mouse and human early embryo. Nature Cell Biology. 2018;20(8):878–87. doi: 10.1038/s41556-018-0144-x.

9. Bhamidipati A, Lewis SA, Cowan NJ. ADP ribosylation factor-like protein 2 (Arl2) regulates the interaction of tubulin-folding cofactor D with native tubulin. J Cell Biol. 2000;149(5):1087–96. doi: 10.1083/jcb.149.5.1087. PubMed PMID: 10831612; PubMed Central PMCID: PMCPMC2174823.

10. Marchetto MC, Belinson H, Tian Y, Freitas BC, Fu C, Vadodaria KC, et al. Altered proliferation and networks in neural cells derived from idiopathic autistic individuals. Molecular Psychiatry. 2017;22(6):820–35. doi: 10.1038/mp.2016.95.

11. Wang M, Wei PC, Lim CK, Gallina IS, Marshall S, Marchetto MC, et al. Increased Neural Progenitor Proliferation in a hiPSC Model of Autism Induces Replication Stress-Associated Genome Instability. Cell Stem Cell. 2020;26(2):221–33.e6. Epub 20200130. doi: 10.1016/j.stem.2019.12.013. PubMed PMID: 32004479; PubMed Central PMCID: PMCPMC7175642.

12. Prem S, Millonig JH, DiCicco-Bloom E. Dysregulation of Neurite Outgrowth and Cell Migration in Autism and Other Neurodevelopmental Disorders. In: DiCicco-Bloom E, Millonig JH, editors. Neurodevelopmental Disorders: Employing iPSC Technologies to Define and Treat Childhood Brain Diseases. Cham: Springer International Publishing; 2020. p. 109–53.

13. Kamiya A, Tan PL, Kubo K, Engelhard C, Ishizuka K, Kubo A, et al. Recruitment of PCM1 to the centrosome by the cooperative action of DISC1 and BBS4: a candidate for psychiatric illnesses. Arch Gen Psychiatry. 2008;65(9):996–1006. doi: 10.1001/archpsyc.65.9.996. PubMed PMID: 18762586; PubMed Central PMCID: PMCPMC2727928.

14. Weghe JCVD, Gomez A, Doherty D. The Joubert–Meckel–Nephronophthisis Spectrum of Ciliopathies. Annual Review of Genomics and Human Genetics. 2022;23(1):301–29. doi: 10.1146/annurev-genom-121321-093528. PubMed PMID: 35655331.

15. Zhang W, Kim PJ, Chen Z, Lokman H, Qiu L, Zhang K, et al. MiRNA-128 regulates the proliferation and neurogenesis of neural precursors by targeting PCM1 in the developing cortex. eLife. 2016;5:e11324. doi: 10.7554/eLife.11324.

16. Buchman JJ, Tseng H-C, Zhou Y, Frank CL, Xie Z, Tsai L-H. Cdk5rap2 Interacts with Pericentrin to Maintain the Neural Progenitor Pool in the Developing Neocortex. Neuron. 2010;66(3):386–402. doi: 10.1016/j.neuron.2010.03.036.

17. Zaqout S, Blaesius K, Wu YJ, Ott S, Kraemer N, Becker LL, et al. Altered inhibition and excitation in neocortical circuits in congenital microcephaly. Neurobiol Dis. 2019;129:130–43. Epub 20190515. doi: 10.1016/j.nbd.2019.05.008. PubMed PMID: 31102767.

18. Lancaster MA, Renner M, Martin C-A, Wenzel D, Bicknell LS, Hurles ME, et al. Cerebral organoids model human brain development and microcephaly. Nature. 2013;501(7467):373-9. doi: 10.1038/nature12517.

19. Bond J, Roberts E, Springell K, Lizarraga S, Scott S, Higgins J, et al. A centrosomal mechanism involving CDK5RAP2 and CENPJ controls brain size. Nature Genetics. 2005;37(4):353–5. doi: 10.1038/ng1539.

20. Zhou C, Cunningham L, Marcus AI, Li Y, Kahn RA. Arl2 and Arl3 regulate different microtubule-dependent processes. Mol Biol Cell. 2006;17(5):2476–87. Epub 20060308. doi: 10.1091/mbc.e05-10-0929. PubMed PMID: 16525022; PubMed Central PMCID: PMCPMC1446103.

21. Francis JW, Turn RE, Newman LE, Schiavon C, Kahn RA. Higher order signaling: ARL2 as regulator of both mitochondrial fusion and microtubule dynamics allows integration of 2 essential cell functions. Small GTPases. 2016;7(4):188–96. Epub 20160711. doi: 10.1080/21541248.2016.1211069. PubMed PMID: 27400436; PubMed Central PMCID: PMCPMC5129891.

22. Chen K, Koe CT, Xing ZB, Tian X, Rossi F, Wang C, et al. Arl2-and Msps-dependent microtubule growth governs asymmetric division. J Cell Biol. 2016;212(6):661–76. Epub 20160307. doi: 10.1083/jcb.201503047. PubMed PMID: 26953351; PubMed Central PMCID: PMCPMC4792071.

23. Shern JF, Sharer JD, Pallas DC, Bartolini F, Cowan NJ, Reed MS, et al. Cytosolic Arl2 is complexed with cofactor D and protein phosphatase 2A. J Biol Chem. 2003;278(42):40829–36. Epub 20030811. doi: 10.1074/jbc.M308678200. PubMed PMID: 12912990.

24. Kahn RA, Volpicelli-Daley L, Bowzard B, Shrivastava-Ranjan P, Li Y, Zhou C, et al. Arf family GTPases: roles in membrane traffic and microtubule dynamics. Biochemical Society Transactions. 2005;33(6):1269–72. doi: 10.1042/bst0331269.

25. Nithianantham S, Le S, Seto E, Jia W, Leary J, Corbett KD, et al. Tubulin cofactors and Arl2 are cage-like chaperones that regulate the soluble αβ-tubulin pool for microtubule dynamics. Elife. 2015;4. Epub 20150724. doi: 10.7554/eLife.08811. PubMed PMID: 26208336; PubMed Central PMCID: PMCPMC4574351.

26. Newman LE, Schiavon CR, Turn RE, Kahn RA. The ARL2 GTPase regulates mitochondrial fusion from the intermembrane space. Cell Logist. 2017;7(3):e1340104. Epub 20170623. doi: 10.1080/21592799.2017.1340104. PubMed PMID: 28944094; PubMed Central PMCID: PMCPMC5602422.

27. Schiavon CR, Turn RE, Newman LE, Kahn RA. ELMOD2 regulates mitochondrial fusion in a mitofusin-dependent manner, downstream of ARL2. Mol Biol Cell. 2019;30(10):1198–213. Epub 20190313. doi: 10.1091/mbc.E18-12-0804. PubMed PMID: 30865555; PubMed Central PMCID: PMCPMC6724520.

28. Davidson AE, Schwarz N, Zelinger L, Stern-Schneider G, Shoemark A, Spitzbarth B, et al. Mutations in ARL2BP, encoding ADP-ribosylation-factor-like 2 binding protein, cause autosomal-recessive retinitis pigmentosa. Am J Hum Genet. 2013;93(2):321–9. Epub 20130711. doi: 10.1016/j.ajhg.2013.06.003. PubMed PMID: 23849777; PubMed Central PMCID: PMCPMC3738823.

29. Cai XB, Wu KC, Zhang X, Lv JN, Jin GH, Xiang L, et al. Whole-exome sequencing identified ARL2 as a novel candidate gene for MRCS (microcornea, rod-cone dystrophy, cataract, and posterior staphyloma) syndrome. Clinical Genetics. 2019;96(1):61–71.

30. Sharer JD, Shern JF, Van Valkenburgh H, Wallace DC, Kahn RA. ARL2 and BART enter mitochondria and bind the adenine nucleotide transporter. Mol Biol Cell. 2002;13(1):71–83. doi: 10.1091/mbc.01-05-0245. PubMed PMID: 11809823; PubMed Central PMCID: PMCPMC65073.

31. Geraldo S, Khanzada UK, Parsons M, Chilton JK, Gordon-Weeks PR. Targeting of the F-actin-binding protein drebrin by the microtubule plus-tip protein EB3 is required for neuritogenesis. Nature Cell Biology. 2008;10(10):1181–9. doi: 10.1038/ncb1778.

32. Newman LE, Zhou CJ, Mudigonda S, Mattheyses AL, Paradies E, Marobbio CM, et al. The ARL2 GTPase is required for mitochondrial morphology, motility, and maintenance of ATP levels. PLoS One. 2014;9(6):e99270. Epub 20140609. doi: 10.1371/journal.pone.0099270. PubMed PMID: 24911211; PubMed Central PMCID: PMCPMC4050054.

33. Fong KW, Choi YK, Rattner JB, Qi RZ. CDK5RAP2 is a pericentriolar protein that functions in centrosomal attachment of the gamma-tubulin ring complex. Mol Biol Cell. 2008;19(1):115–25. Epub 20071024. doi: 10.1091/mbc.e07-04-0371. PubMed PMID: 17959831; PubMed Central PMCID: PMCPMC2174194.

34. Zhu W, Shenoy A, Kundrotas P, Elofsson A. Evaluation of AlphaFold-Multimer prediction on multi-chain protein complexes. Bioinformatics. 2023;39(7). doi: 10.1093/bioinformatics/btad424.

35. Yu D, Chojnowski G, Rosenthal M, Kosinski J. AlphaPulldown-a python package for protein-protein interaction screens using AlphaFold-Multimer. Bioinformatics. 2023;39(1). doi: 10.1093/bioinformatics/btac749. PubMed PMID: 36413069; PubMed Central PMCID: PMCPMC9805587.

36. Mirdita M, Schütze K, Moriwaki Y, Heo L, Ovchinnikov S, Steinegger M. ColabFold: making protein folding accessible to all. Nature Methods. 2022;19(6):679–82. doi: 10.1038/s41592-022-01488-1.

37. Fredriksson S, Gullberg M, Jarvius J, Olsson C, Pietras K, Gustafsdottir SM, et al. Protein detection using proximity-dependent DNA ligation assays. Nat Biotechnol. 2002;20(5):473–7. doi: 10.1038/nbt0502-473. PubMed PMID: 11981560.

38. Uzquiano A, Gladwyn-Ng I, Nguyen L, Reiner O, Götz M, Matsuzaki F, et al. Cortical progenitor biology: key features mediating proliferation versus differentiation. J Neurochem. 2018;146(5):500–25. Epub 20180627. doi: 10.1111/jnc.14338. PubMed PMID: 29570795.

39. Konno D, Shioi G, Shitamukai A, Mori A, Kiyonari H, Miyata T, et al. Neuroepithelial progenitors undergo LGN-dependent planar divisions to maintain self-renewability during mammalian neurogenesis. Nat Cell Biol. 2008;10(1):93–101. Epub 20071216. doi: 10.1038/ncb1673. PubMed PMID: 18084280.

40. Wright ZC, Loskutov Y, Murphy D, Stoilov P, Pugacheva E, Goldberg AFX, et al. ADP-Ribosylation Factor-Like 2 (ARL2) regulates cilia stability and development of outer segments in rod photoreceptor neurons. Sci Rep. 2018;8(1):16967. Epub 20181116. doi: 10.1038/s41598-018-35395-3. PubMed PMID: 30446707; PubMed Central PMCID: PMCPMC6240099.

41. Mitchell-Dick A, Chalem A, Pilaz LJ, Silver DL. Acute Lengthening of Progenitor Mitosis Influences Progeny Fate during Cortical Development in vivo. Dev Neurosci. 2019;41(5-6):300–17. Epub 20200615. doi: 10.1159/000507113. PubMed PMID: 32541147; PubMed Central PMCID: PMCPMC7388066.

42. Zhou Y, Jiang H, Gu J, Tang Y, Shen N, Jin Y. MicroRNA-195 targets ADP-ribosylation factor-like protein 2 to induce apoptosis in human embryonic stem cell-derived neural progenitor cells. Cell Death & Disease. 2013;4(6):e695-e. doi: 10.1038/cddis.2013.195.

43. Iwata R, Casimir P, Vanderhaeghen P. Mitochondrial dynamics in postmitotic cells regulate neurogenesis. Science. 2020;369(6505):858-62. doi: 10.1126/science.aba9760. PubMed PMID: 32792401.

44. Lizarraga SB, Margossian SP, Harris MH, Campagna DR, Han AP, Blevins S, et al. Cdk5rap2 regulates centrosome function and chromosome segregation in neuronal progenitors. Development. 2010;137(11):1907–17. doi: 10.1242/dev.040410. PubMed PMID: 20460369; PubMed Central PMCID: PMCPMC2867323.

45. Paridaen JT, Wilsch-Bräuninger M, Huttner WB. Asymmetric inheritance of centrosome-associated primary cilium membrane directs ciliogenesis after cell division. Cell. 2013;155(2):333–44. doi: 10.1016/j.cell.2013.08.060. PubMed PMID: 24120134.

46. Schmechel DE, Rakic P. A Golgi study of radial glial cells in developing monkey telencephalon: morphogenesis and transformation into astrocytes. Anat Embryol (Berl). 1979;156(2):115–52. doi: 10.1007/bf00300010. PubMed PMID: 111580.

47. Coquand L, Victoria GS, Tata A, Carpentieri JA, Brault JB, Guimiot F, et al. CAMSAPs organize an acentrosomal microtubule network from basal varicosities in radial glial cells. J Cell Biol. 2021;220(8). Epub 20210521. doi: 10.1083/jcb.202003151. PubMed PMID: 34019079; PubMed Central PMCID: PMCPMC8144914.

48. Issa L, Mueller K, Seufert K, Kraemer N, Rosenkotter H, Ninnemann O, et al. Clinical and cellular features in patients with primary autosomal recessive microcephaly and a novel CDK5RAP2 mutation. Orphanet Journal of Rare Diseases. 2013;8(1):59. doi: 10.1186/1750-1172-8-59.

49. Cai XB, Wu KC, Zhang X, Lv JN, Jin GH, Xiang L, et al. Whole-exome sequencing identified ARL2 as a novel candidate gene for MRCS (microcornea, rod-cone dystrophy, cataract, and posterior staphyloma) syndrome. Clin Genet. 2019;96(1):61–71. Epub 20190422. doi: 10.1111/cge.13541. PubMed PMID: 30945270.

50. Gerstner CD, Reed M, Dahl TM, Ying G, Frederick JM, Baehr W. Arf-like Protein 2 (ARL2) Controls Microtubule Neogenesis during Early Postnatal Photoreceptor Development. Cells. 2022;12(1). Epub 20221230. doi: 10.3390/cells12010147. PubMed PMID: 36611941; PubMed Central PMCID: PMCPMC9818799.

51. Zaqout S, Ravindran E, Stoltenburg-Didinger G, Kaindl AM. Congenital microcephaly-linked CDK5RAP2 affects eye development. Ann Hum Genet. 2020;84(1):87–91. Epub 20190729. doi: 10.1111/ahg.12343. PubMed PMID: 31355417.

52. Lois C, Hong EJ, Pease S, Brown EJ, Baltimore D. Germline transmission and tissue-specific expression of transgenes delivered by lentiviral vectors. Science. 2002;295(5556):868-72. Epub 20020110. doi: 10.1126/science.1067081. PubMed PMID: 11786607.

53. Rompani SB, Cepko CL. Retinal progenitor cells can produce restricted subsets of horizontal cells. Proceedings of the National Academy of Sciences. 2008;105(1):192–7. doi: doi:10.1073/pnas.0709979104.

54. Shyu YJ, Liu H, Deng X, Hu CD. Identification of new fluorescent protein fragments for bimolecular fluorescence complementation analysis under physiological conditions. Biotechniques. 2006;40(1):61–6. doi: 10.2144/000112036. PubMed PMID: 16454041.

55. Kodama Y, Hu CD. An improved bimolecular fluorescence complementation assay with a high signal-to-noise ratio. Biotechniques. 2010;49(5):793–805. doi: 10.2144/000113519. PubMed PMID: 21091444.

56. Ma D, Yoon S-I, Yang C-H, Marcy G, Zhao N, Leong W-Y, et al. Rescue of Methyl-CpG Binding Protein 2 Dysfunction-induced Defects in Newborn Neurons by Pentobarbital. Neurotherapeutics. 2015;12(2):477–90. doi: 10.1007/s13311-015-0343-0.

57. Meyer-Dilhet G, Courchet J. In Utero Cortical Electroporation of Plasmids in the Mouse Embryo. STAR Protoc. 2020;1(1):100027. Epub 20200619. doi: 10.1016/j.xpro.2020.100027. PubMed PMID: 32685931; PubMed Central PMCID: PMCPMC7357676.

58. Gujar MR, Gao Y, Teng X, Deng Q, Lin KY, Tan YS, et al. Golgi-dependent reactivation and regeneration of Drosophila quiescent neural stem cells. Dev Cell. 2023;58(19):1933–49 e5. Epub 20230810. doi: 10.1016/j.devcel.2023.07.010. PubMed PMID: 37567172.

59. Evans R, O’Neill M, Pritzel A, Antropova N, Senior A, Green T, et al. Protein complex prediction with AlphaFold-Multimer. bioRxiv. 2022:2021.10.04.463034. doi: 10.1101/2021.10.04.463034.

60. Deng Q, Wang C, Koe CT, Heinen JP, Tan YS, Li S, et al. Parafibromin governs cell polarity and centrosome assembly in Drosophila neural stem cells. PLoS Biol. 2022;20(10):e3001834. Epub 20221012. doi: 10.1371/journal.pbio.3001834. PubMed PMID: 36223339; PubMed Central PMCID: PMCPMC9555638.

61. Jakobs MAH, Dimitracopoulos A, Franze K. KymoButler, a deep learning software for automated kymograph analysis. eLife. 2019;8:e42288. doi: 10.7554/eLife.42288.

